# The probability of evolutionary rescue with competition: scaling-up from a single species to a community

**DOI:** 10.64898/2026.09.11.751028

**Authors:** Kuangyi Xu, Matthew M. Osmond

## Abstract

When ecological communities experience environmental change, continued coexistence may require rapid adaptation to avoid extinction, i.e., evolutionary rescue. This rescue can be inhibited by competition. Combining Lotka-Volterra and population genetic models, we investigate how both within- and between-species competition impact the probability of rescue. With a single species we derive closed-form approximations that quantify the negative impact of competition. We then use a two-species model to examine how the sweep of a rescue mutant in one species inhibits establishment of rescue mutants in another, a process we term survival interference. This interference is strongest when the stronger competitor begins recovering first, creating eco-evolutionary priority effects despite conditions permitting deterministic coexistence. Finally, considering larger competitive communities, we document how survival interference reduces the number of rescued species but weakens with species richness and unevenness. These results move us closer to predicting evolutionary rescue in ecological communities.

## 1 Introduction

When an ecological community is hit by a large environmental change, such as a coral reef under thermal stress or a gut microbiome exposed to a high dose of antibiotics, multiple species may find themselves outside their niche and therefore in demographic decline. Post-change coexistence and biodiversity then depend not only on altered interaction strengths (Letten et al., 2021) but also on the chance that species adapt fast enough to be rescued from extinction by evolution (Gomulkiewicz and Holt, 1995; Bell, 2017). “Community rescue” experiments, such as those that have exposed large microbial (Low-Décarie et al., 2015) and phytoplankton (Fugère et al., 2020) communities to herbicides, are beginning to document large shifts in community composition and species diversity after dramatic environmental change. This is not simply independent evolutionary rescue in each species. For example, the evolution of drug resistance in one bacterial species has been shown to repeatedly exclude others (Kivikoski et al., 2026). To understand and predict these community-level changes we need a theoretical framework that merges ecological interactions and evolutionary genetics and yet is simple enough to scale up to many species.

Rescue of a focal species is critically influenced by competition. Before considering multiple species, it is first helpful to understand the effect of within-species competition. Models of rescue via the evolution of quantitative traits (e.g., Gomulkiewicz and Holt, 1995), where genetic variation is abundant and rescue depends on the (roughly deterministic) rate of evolution, show that intraspecific competition is expected to inhibit rescue by reducing population size (Chevin and Lande, 2010) and genetic variation (Nordstrom et al., 2023). Other models assume rescue occurs via the stochastic establishment of rare adaptive alleles at major-effect loci (e.g., Orr and Unckless, 2008). These studies emphasize that intraspecific competition changes how the probability of rescue depends on the severity of stress, with important implications for the evolution of drug resistance (Read et al., 2011). In particular, a faster decline of the wildtype (e.g., due to a higher drug dose) can increase the probability of rescue by releasing existing rescue mutants from competition (Uecker et al., 2014; Wilson et al., 2017; Day and Read, 2016; Czuppon et al., 2023).

Competition between species also affects rescue. Quantitative genetics models show that the presence of a competitor generally inhibits rescue of a focal species (de Mazancourt et al., 2008; Johansson, 2008), unless it imposes selection that sufficiently hastens adaptation (Jones, 2008; Osmond and de Mazancourt, 2013). The effect of interspecific competition on rescue via the stochastic establishment of large effect mutations is less clear. Individual-based simulations of a community of competitors, equal in all respects except initial abundance and that each require the fixation of a new mutation for survival, show that initially larger populations are more likely to be rescued and their rescue can inhibit the rescue of others (Van Eldijk et al., 2020). We are interested in exploring this inhibition in more detail. In particular, we suspect that the sweep of a rescue mutant in one species will inhibit establishment of a rescue mutant in another species. We term this mutual inhibition of rescue “survival interference”, by analogy to clonal interference, where beneficial mutations compete for fixation within a population and reduce genetic diversity (Otto, 2021). By hindering coexistence, we expect survival interference will be a key determinant of post-change community structure and diversity.

Our broad goal here is to generate more intuition about the effect of competition on the probability of rescue and to develop a framework that can be extended to communities with many competing species. To do so, we build relatively simple models and derive analytical and numerical approximations. We first investigate the impact of intraspecific competition in a single-species system, deriving new closed-form approximations for the probability of rescue. We then extend the analysis to a two-species system, where we can examine survival interference in detail. Finally, we show how this approach can be scaled-up to a larger communities of competing species.

## 2 Methods

Our aim is to build a conceptually simple and tractable stochastic model of evolutionary rescue with competition. We consider haploid populations with non-overlapping generations that experience an abrupt environmental change. The number of offspring produced by an individual with fitness *W* is Poisson with mean *W* . Mutations then occur. Fitness is determined by one major-effect locus. Initially, the population is primarily wildtype (with *W <* 1) and suffers demographic decline, but may be rescued through the establishment of a rare rescue allele that either pre-existed before the environmental shift in the standing genetic variation or arises during demographic decline by *de novo* mutation. Disentangling whether adaptation occurs via standing genetic variation (SGV) or via *de novo* mutations (DNM) has long been a central question in evolutionary biology (Barrett and Schluter, 2008; Bomblies and Peichel, 2022; Kersten et al., 2023), with important implications for resistance management (Délye et al., 2013; Hawkins et al., 2019). We therefore distinguish between rescue arising from SGV and DNM and compare their relative contributions.

### 2.1 One species

Consider a locus with two alleles, the wildtype *a* and the mutant *A*. Let *n*_*i*_ be the density of allele *i* and *n*_*i*_*K* the absolute number, where *K* is a scaling parameter (e.g., habitat area). The Malthusian fitnesses of the two alleles under the *r* − *α* version of the Lotka-Volterra model (Mallet, 2012) are 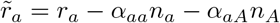 and 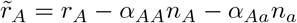, where *r*_*i*_ is the intrinsic growth rate of allele *i* and *α*_*ij*_ is the strength of competition of allele *j* on allele *i*. We restrict to *α*_*ij*_ ≥ 0 to focus on competition but the same model could describe cooperation. Throughout we set *α*_*ii*_ = 1 for simplicity; this can be done without loss of generality by considering the density of allele *i* scaled by *α*_*ii*_.

The number of offspring with allele *i* in generation *t* + 1 is Poisson distributed with mean 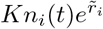. Mutations then determine the number of adults of each type. We assume allele *a* mutates to allele *A* with probability *µ* and ignore back mutations. A continuous-time approximation of the dynamics of the expected densities is

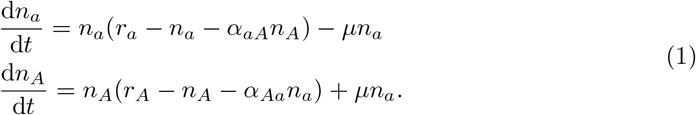

Note that the mutant population is initially extremely small and may therefore be lost due to demographic stochasticity despite having a positive growth rate. Consequently, evolutionary rescue is not guaranteed and depends on whether a mutant lineage successfully establishes.

The same model structure describes the dynamics before and after the environmental change, which only changes parameter values (an asterisk denotes parameter values before the environmental shift). Before the environmental shift, we assume the wildtype can grow when rare, 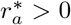, and has a higher intrinsic growth rate than the mutant, 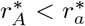. After the environmental shift, the wildtype cannot grow but the mutant can, *r*_*a*_ *<* 0 *< r*_*A*_. Both before and after environmental change, mutants are assumed to arise infrequently, *µ*^∗^, *µ* ≪ 1.

### 2.2 Two species

We then consider two species, labeled 1 and 2. Species 1 has a locus with alleles *a* and *A* and species 2 has a locus with alleles *b* and *B*. Alleles *a* and *b* are wildtype and alleles *A* and *B* are mutants. The absolute number of alleles are *n*_*a*_*K*_1_, *n*_*A*_*K*_1_, *n*_*b*_*K*_2_ and *n*_*B*_*K*_2_, where *n*_*i*_ is the density of allele *i* and *K*_*j*_ is a scaling parameter for species *j*. For simplicity, we assume that all intraspecific competition coefficients are one and explore the effect of interspecific competition from species *j* on species *i*, with competition coefficient *β*_*ij*_. Malthusian fitness of allele *a* in species 1 is then 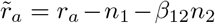, where *n*_1_ = *n*_*a*_ +*n*_*A*_ and *n*_2_ = *n*_*b*_ +*n*_*B*_ are the densities of species 1 and 2, respectively. The fitnesses of the other alleles are written analogously.

The number of offspring in species *j* with allele *i* in generation *t* +1 is Poisson with mean 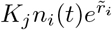. Offspring with allele *a* mutate to allele *A* with probability *µ*_1_ and offspring with allele *b* mutate to allele *B* with probability *µ*_2_. A continuous-time approximation for the dynamics of the expected densities is

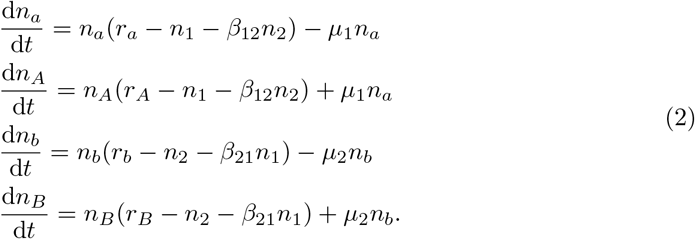

As in the single-species case, the mutants in both species are initially rare and thus may be lost by demographic stochasticity despite a potentially positive growth rate.

As in the one species case, we assume weak mutation (*µ*^∗^, *µ* ≪ 1) and wildtype advantage before the environmental change (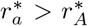 and 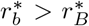) so that the mutants are initially rare. We assume that the two species stably coexist prior to the environmental change. Defining niche overlap 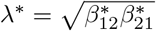 and competitive ability ratio 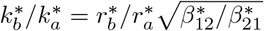 (Letten et al., 2021), the wildtypes can stably coexist when 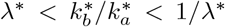. This requires that their niches do not completely overlap, *λ*^∗^ *<* 1. The equilibrium densities of species 1 and 2 are 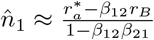 and 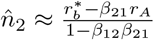, respectively.

### 2.3 Simulations

For a given set of parameter values, we obtain the rescue probability by simulating 10^5^ replicates (in R 4.3.2). For rescue from standing genetic variation, we evolved the population for 6 times the initial population size to attain mutation-selection-drift balance prior to the environmental shift. We consider a species rescued when the number of mutant alleles exceeds 500 and the product of the realized growth rate of the mutant and the number of mutant alleles exceeds 2.

## 3 Results

### 3.1 One species

#### 3.1.1 Mutant establishment

In Section S1.1 we combine the establishment probability derived by Uecker and Hermisson, 2011 with deterministic wildtype decline and interpolate over the strength of competition (from *α*_*Aa*_ = 0 to *α*_*Aa*_ = 1) to approximate the probability a mutant arising at time *t* establishes,

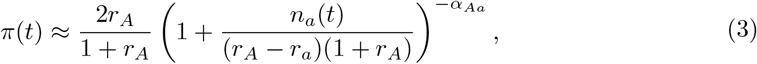

where

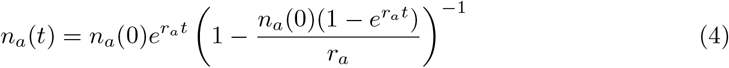

is the density of the wildtype *a* at time *t*. This approximation works well even for *α*_*Aa*_ *>* 1 (Figure 1b) and gives immediate intuition. In particular, when the mutant has a low intrinsic growth rate, *r*_*A*_ ≪ 1, the effect of competition on establishment is governed by the density of the wildtype at the time the mutant arises relative to the intrinsic mutant advantage, *n*_*a*_(*t*)*/*(*r*_*A*_ − *r*_*a*_), consistent with Czuppon et al., 2023, who focused on mutants in the standing variation (*t* = 0). Faster decline of the wildtype reduces intraspecific competition (competitive release) and therefore increases the establishment probability of mutants arising at a given time. As time proceeds, the wildtype disappears (*n*_*a*_(*t*) → 0) and the establishment probability converges to 2*r*_*A*_*/*(1 + *r*_*A*_).

**Figure 1:**
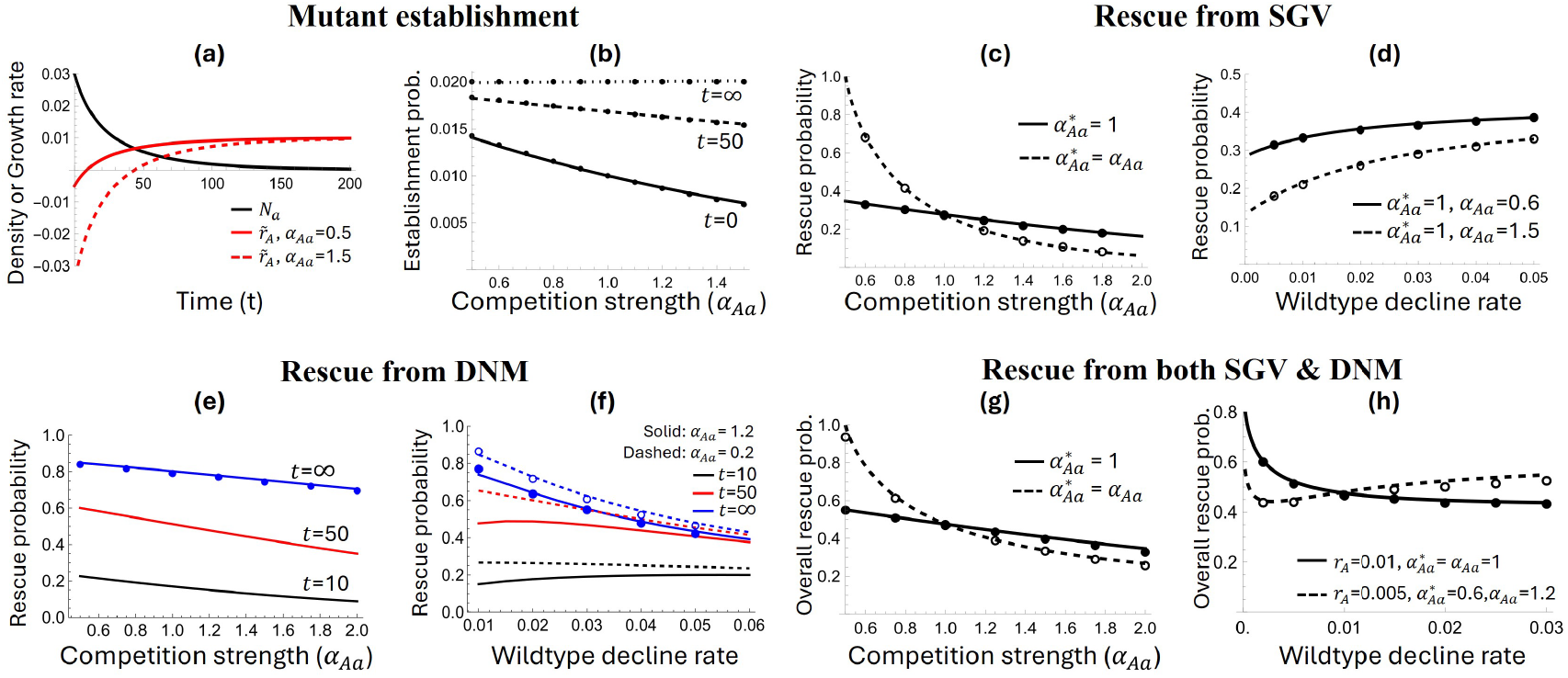
Effects of intraspecific competition and wildtype decline rate on rescue of an isolated species. **(a)** Dynamics of wildtype density and the resulting growth rate of a rare rescue mutant, 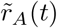. **(b)** Mutant establishment probability, *π*(*t*), as a function of per capita competition strength from the wildtype, *α*_*Aa*_. Lines are the closed-form approximation (Equation 3) and dots are the more accurate Equation S3. **(c)-(h)** Impacts of competition strength and wildtype decline rate on the probability of rescue from SGV, DNM, or both. Lines are analytical predictions (Equations 5, 6, and S9) and dots are from simulations. Parameters in (a) & (b): *n*_*a*_(0) = 0.03, *r*_*a*_ = −0.02, *r*_*A*_ = 0.01. Unless otherwise specified, parameters in (c)-(h): 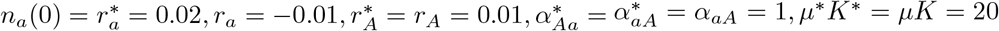, except for (e) and (f) where *µ*^∗^*K*^∗^ = *µK* = 100 to keep rescue probabilities at intermediate levels for clearer illustration.

#### 3.1.2 Rescue from standing genetic variation (SGV)

Assuming mutation-selection-drift balance before the environmental shift and that the probability that a mutant in the SGV establishes is small, the probability of rescue from the SGV is (Section S1.2)

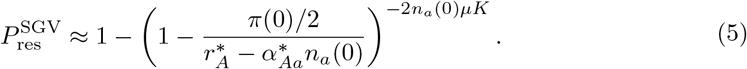

Competition affects rescue from SGV in two ways. First, the probability of rescue decreases with the strength of intraspecific competition in the new environment (solid lines in Figure 1c), *α*_*Aa*_, due to reduced establishment probability, *π*(0), as mentioned above. Rescue by SGV also increases with a higher realized growth rate of mutants prior to the environmental change, 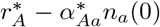, which decreases with the strength of pre-shift intraspecific competition, 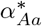. Increasing the strength of intraspecific competition before and after the environmental shift therefore causes the probability of rescue to decline rapidly (dashed lines in Figure 1c), due to both a lower initial number of mutants and a lower establishment probability.

The probability of rescue from SGV increases with a faster decline of the wildtype (Figure 1d), owing to a higher establishment probability of rescue mutants. This competitive release is more pronounced when competition is strong (compare the rate of increase in the dashed and solid lines in Figure 1d) since the wildtype then suppresses mutant establishment more.

#### 3.1.3 Rescue from *de novo* mutations (DNM)

We now consider rescue from mutations arising during the wildtype’s decline. Again interpolating between *α*_*Aa*_ = 0 and *α*_*Aa*_ = 1, a closed-form approximation for the probability of rescue from DNM is (Section S1.3)

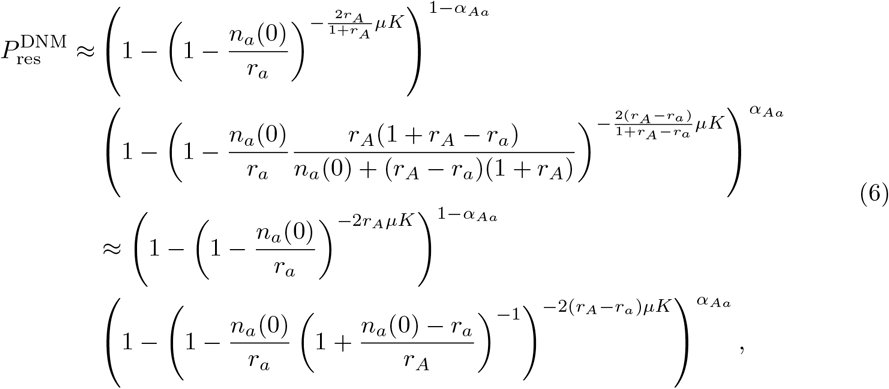

where the second approximation assumes *r*_*a*_, *r*_*A*_ ≪ 1. Setting *α*_*Aa*_ = 0 shows that without competition from the wildtype, the probability of rescue depends on the product of the cumulative number of wildtypes, ln(1−*n*_*a*_(0)*/r*_*a*_), and the rate at which establishing mutants are produced, 2*r*_*A*_*µK*. Comparing *α*_*Aa*_ = 0 and *α*_*Aa*_ = 1, we see that competition from the wildtype can be interpreted as effectively increasing the establishment probability from the typical 2*r*_*A*_, twice the mutant’s growth rate, to 2(*r*_*A*_ − *r*_*a*_), twice the mutant’s selective advantage, and modulating the cumulative number of wildtypes in a way that depends only on (*n*_*a*_(0) − *r*_*a*_)*/r*_*A*_. The larger the initial wildtype population, *n*_*a*_(0), the more competition lowers the probability of rescue.

The probability of rescue from DNM ultimately declines with increased competition from the wildtype and with wildtype decline rate (blues lines in Figure 1e,f). However, by deriving the probability of rescue by time *t* (Equation S9), we see that in the short term the probability of rescue peaks at an intermediate wildtype decline rate (black and red lines in Figure 1f). Intuitively, increasing the wildtype decline rate can benefit rescue by increasing the establishment probability of mutants via relaxed competition, but also imposes a cost of fewer *de novo* mutants. Stronger competition from the wildtype increases the benefit of competitive release, causing the probability of rescue to peak at higher decline rates (Figure S1). However, as time proceeds, the benefit weakens and the cost strengthens (Figure S2) and ultimately the rescue probability monotonically decreases with faster wildtype decline (blue line in Figure 1f), aligning with Uecker et al., 2014. By increasing the benefit, stronger competition from the wildtype causes the rescue probability to peak at intermediate decline rates for longer (Figure S1).

#### 3.1.4 Rescue from either SGV and DNM

Considering both SGV and DNM, the overall rescue probability is 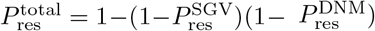. Increasing competition from the wildtype lowers the probability of rescue (Figure 1g) and the relative contribution of SGV (Figure S3a). These patterns are more pronounced when competition before environmental change is also strengthened, as this reduces the number of mutants in the SGV (compare solid and dashed lines in Figure 1g).

As reported by Uecker et al., 2014, the overall rescue probability either decreases monotonically with increasing wildtype decline rate or exhibits a minimum at an intermediate decline rate (Figure 1h). The intermediate minimum occurs when the increase in rescue by SGV outweighs the reduction in rescue by DNM (Figure S3b). This is more likely to happen when the mutant experiences weak competition in the original environment but strong competition in the new environment, increasing the amount of SGV and the benefit of competitive release (dashed line in Figure 1h). While interesting, we find this non-monotonic pattern in a relatively small parameter range.

### 3.2 Two species

#### 3.2.1 Rescue probabilities

With two species there are four possible outcomes after the environmental shift: (1) both species go extinct, (2) species 1 goes extinct and species 2 is rescued, (3) species 1 is rescued and species 2 goes extinct, and (4) both species are rescued and coexist. We denote the probability of each of the four outcomes as *P*_00_, *P*_01_, *P*_10_ and *P*_11_, respectively. It is equivalent to calculate the rescue probabilities of species 1 and 2, *P*_res,1_ = *P*_10_ + *P*_11_ and *P*_res,2_ = *P*_01_ + *P*_11_, and the correlation between rescue events in the two species, 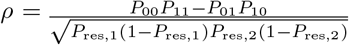. A negative correlation means that the rescue of both species is less likely than expected based on the single species rescue probabilities, *P*_11_ *< P*_res,1_*P*_res,2_, i.e., there is survival interference, and the magnitude of this negative correlation indicates the strength of survival interference. We are interested in how competition affects the rescue probability of each species as well as post-change coexistence. As we will see, factors that increase the rescue probability of each species do not necessarily increase the probability of coexistence.

The dynamics with two species can be divided into two stages, depending on whether a rescue mutant has started to sweep or not. In the first stage, rescue mutants are rare in both species and the dynamics of the two species are mainly determined by competition between the two declining wildtypes (stage I in Figure 2a). The second stage begins when a rescue mutant starts to sweep in one species, which slightly speeds up the decline of both wildtypes (stage II in Figure 2a) and, more importantly, interferes with the establishment of rescue mutants in the other species. If the mutant in species 1 (*A*) does not sweep, the establishment probability of the mutant in species 2 (*B*) increases monotonically over time (solid line in Figure 2b) due to competitive release from both wildtypes (Figure 2c). If *A* does sweep, it reduces the establishment probability of *B* and this reduction is more severe the earlier *A* starts sweeping (compare the dashed and dotted lines in Figure 2b). As shown below, factors that cause mutants to sweep earlier tend to strengthen interference, inhibiting coexistence. Based on this logic, we can approximate the rescue probabilities by conditioning the density dynamics and establishment probabilities on when a mutant begins to sweep and integrate over the distribution of waiting times until a rescue mutant starts sweeping (Section S2.1). These approximations give intuition and speed up numerical evaluations of the rescue probabilities, but, due to the complications of interference between mutants, do not lead to closed-form expressions.

**Figure 2:**
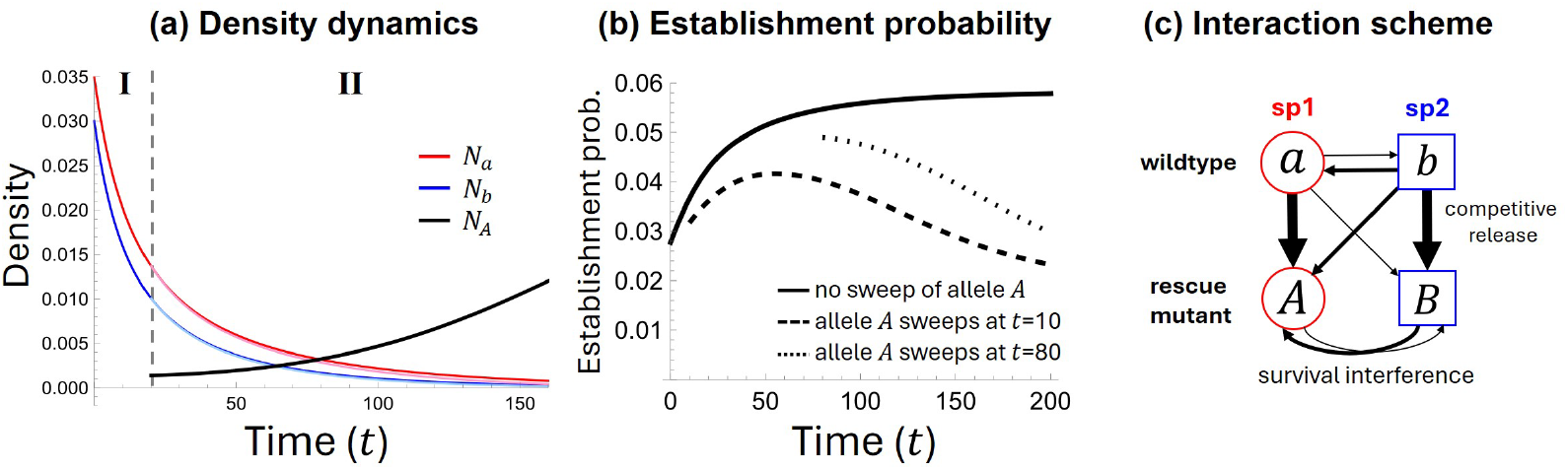
Survival interference between rescue mutations in two species. **(a)** Approximate dynamics of the wildtype in each species (*a* and *b*) without a mutant sweep (dark red and blue lines; Equation S18) and after the rescue mutant in species 1 (*A*) starts sweeping at *t* = 20 (light red and blue lines; strongly overlapping dark lines; Equation S19). **(b)** Establishment probability of the rescue mutant in species 2 (*B*) without the sweep of *A* (solid line) and conditioned on *A* starting to sweep at *t* = 10 and *t* = 80 (broken lines). See Section S2.1 for derivations. **(c)** Schematic of interactions between alleles within and between species. Decline of the wildtypes leads to competitive release of the mutants. The sweep of a mutant in one species interferes with the sweep of a mutant in the other, which we call survival interference. Parameters: *n*_*a*_(0) = 0.035, *n*_*b*_(0) = 0.03, 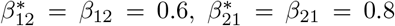, *r*_*A*_ = 0.025, *r*_*B*_ = 0.03, *r*_*a*_ = −0.015, *r*_*b*_ = −0.02, 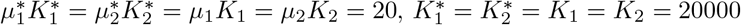.

#### 3.2.2 Impacts of competition strength

Increasing the strength of competition of species 2 on species 1, *β*_12_, inhibits the rescue of species 1 and facilitates the rescue of species 2 (Figure 3a,b). However, the latter effect is generally much weaker (compare slopes in Figure 3a with slopes 3b) because, although increasing *β*_12_ causes the other wildtype, *a*, to decline faster, which would by itself increase establishment of *B*, the faster decline of *a* slows down the decline of *b*, with which *B* competes more strongly.

**Figure 3:**
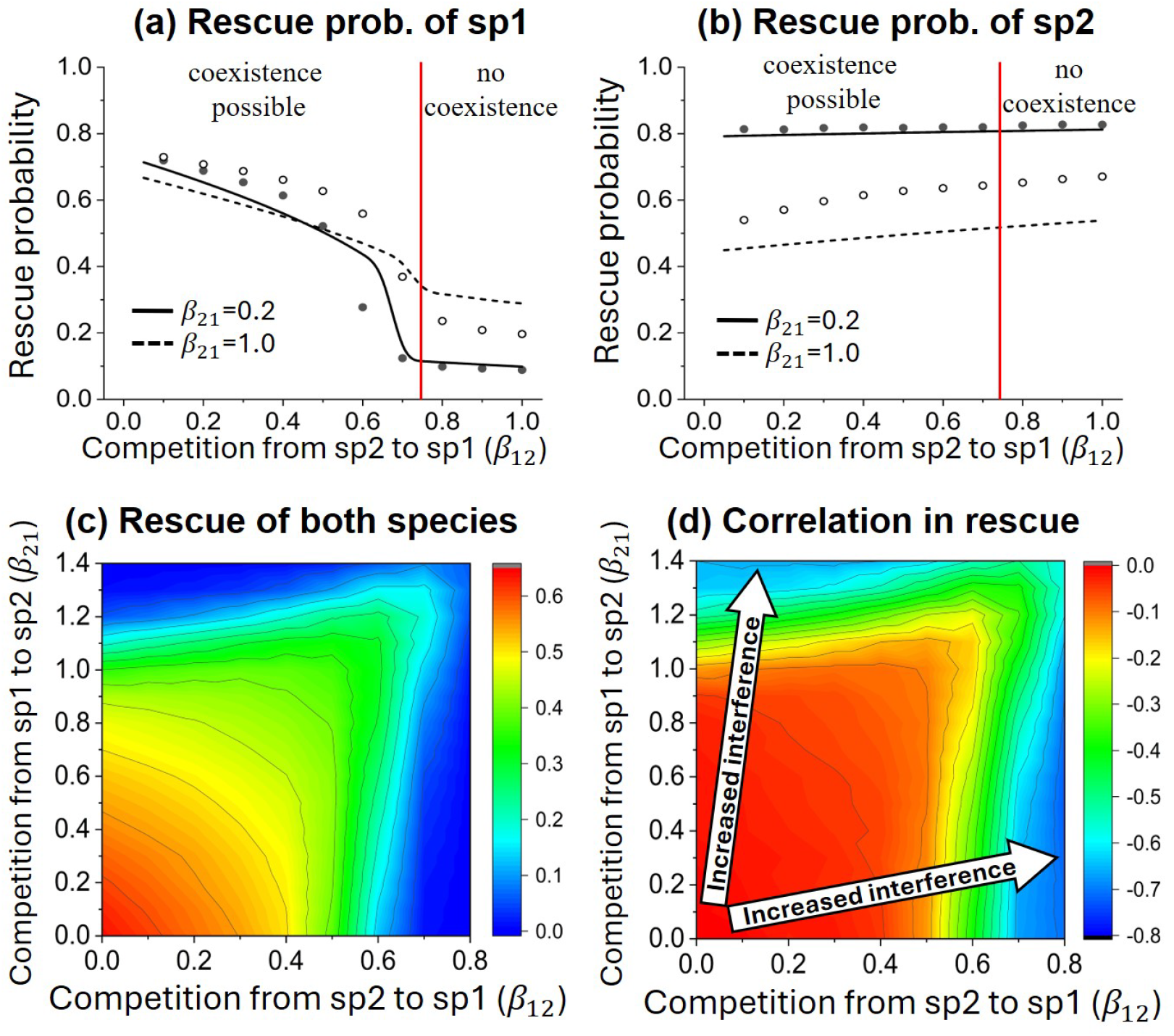
Impact of interspecific competition strength on rescue from DNM with two species. Results are similar for rescue from SGV (Figure S5). Lines in panels (a) and (b) are analytical approximations (Section S2). Dots and contours are from simulations. In panels (a) and (b), coexistence is not possible when *β*_12_ *> r*_*A*_*/r*_*B*_ = 0.75 (indicated by the red vertical line), in which case species 1 will be excluded if the mutant establishes in species 2. Parameters: *r*_*a*_ = *r*_*b*_ = −0.01, *r*_*A*_ = 0.015, *r*_*B*_ = 0.02, *µ*_1_*K*_1_ = *µ*_2_*K*_2_ = 50, *K*_1_ = *K*_2_ = 200000, and the initial density of the two species is fixed as *n*_1,0_ = *n*_2,0_ = 0.03 (results are similar when the initial density at coexistence depends on the competition strength; Figure S6).

Not surprisingly, coexistence is most likely when competition is weak in both directions (bottom-left region in Figure 3c) as the rescue probability of each species is high (Figure 3a,b) and survival interference is minimal (bottom-left region in Figure 3d). For a given total strength of competition, *β*_12_ + *β*_21_, the lowest probability of species coexistence occurs when competition is strongly asymmetric (upper-left and bottom-right corners in Figure 3c). This is due to both strengthened interference (Figure 3d) and a reduced rescue probability of one of the two species (Figure S4). With asymmetry, the sweep of the rescue mutant in the stronger competitor strongly inhibits the establishment of the rescue mutant in the other.

#### 3.2.3 Impacts of wildtype decline rate

As in the one-species case, faster decline of a focal wildtype promotes rescue from SGV (Figure 4a) and inhibits rescue from DNM in the long-term (Figure 5a). These effects are relatively unaffected by competition from the other species (Figures 4a and 5a). On the other hand, increasing the rate of decline of a focal species promotes the rescue of its competitor by both SGV (Figure 4b) and DNM (Figure 5b) and this between-species competitive release is more prominent when competition on the other species is stronger (Figures 4b and 5b).

**Figure 4:**
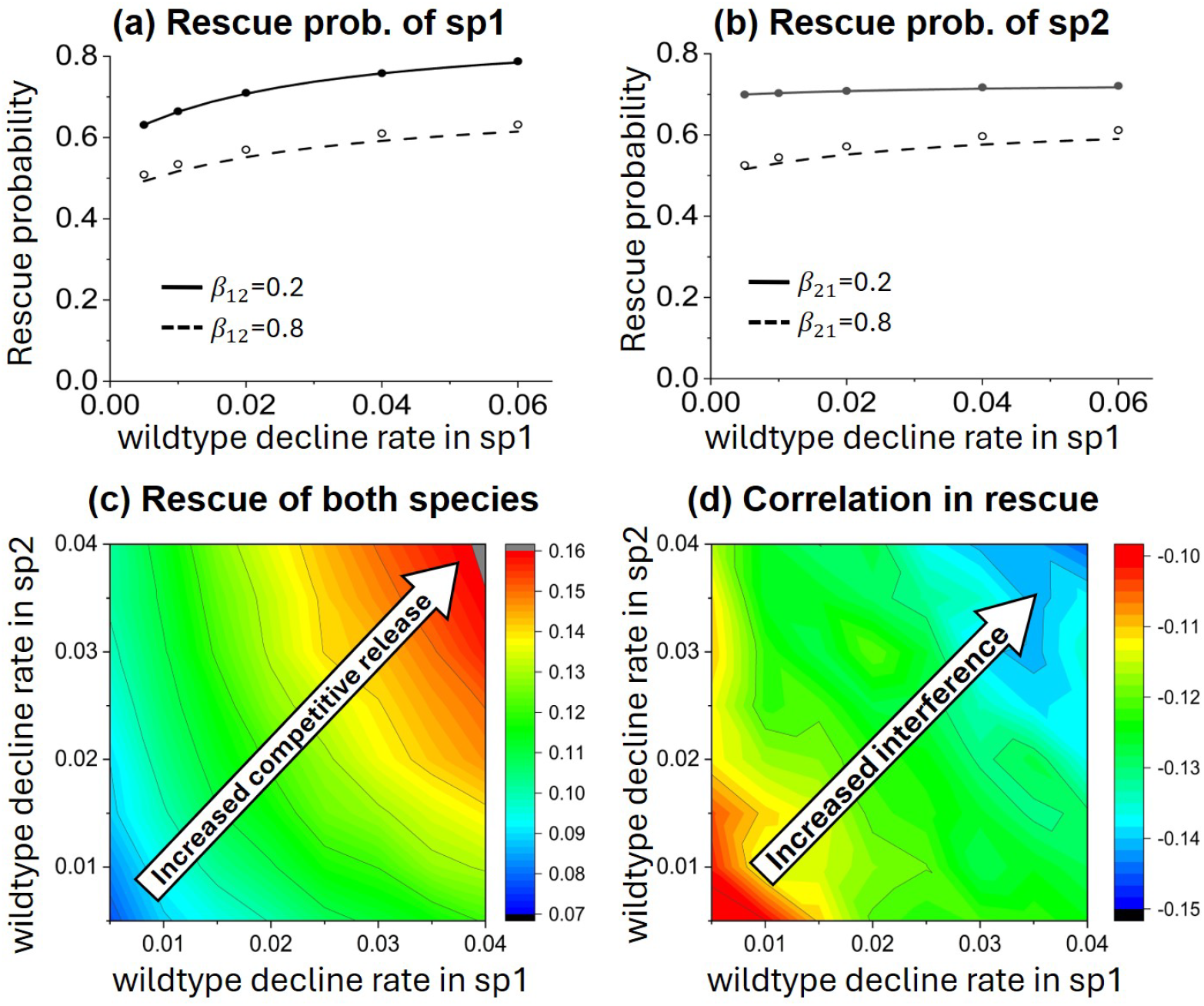
Impact of wildtype decline rates on rescue from SGV with two species. Lines in panels (a) and (b) are analytical approximations (Section S2). Dots and contours are from simulations based on 10^5^ replicates. In panel (a) *β*_21_ = 0.5 and in panel (b) *β*_12_ = 0.5. In panels (a) and (b) *µ*_1_*K*_1_ = *µ*_2_*K*_2_ = 20. In panels (c) and (d) *β*_12_ = 0.2, *β*_21_ = 0.8, *µ*_1_*K*_1_ = *µ*_2_*K*_2_ = 100. Other parameters: 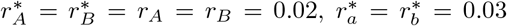,*K*_1_ = *K*_2_ = 200000.

**Figure 5:**
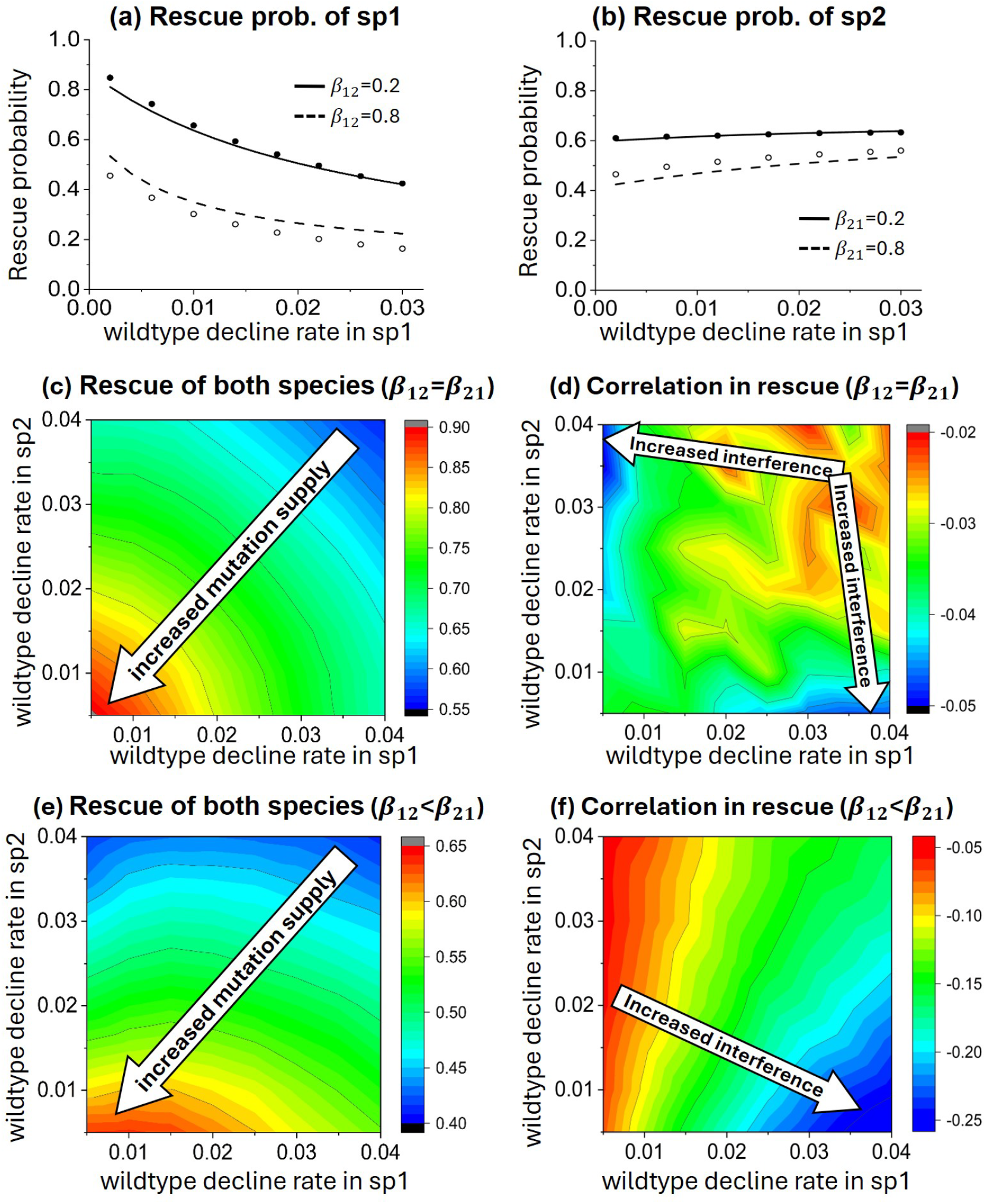
Impact of wildtype decline rates on rescue from DNM with two species. Lines in panels (a) and (b) are analytical approximations (Section S2). Dots and contours are from simulations with 10^5^ replicates. In panel (a) *β*_21_ = 0.8 and in panel (b) *β*_12_ = 0.8. In panels (a) and (b) *r*_*b*_ = −0.02, *µ*_1_*K*_1_ = *µ*_2_*K*_2_ = 50. In panels (c) and (d) *β*_12_ = *β*_21_ = 0.8. In panels (e) and (f) *β*_12_ = 0.2, *β*_21_ = 0.8. Other parameters: *r*_*A*_ = *r*_*B*_ = 0.02, *n*_1,0_ = *n*_2,0_ = 0.03, *µ*_1_*K*_1_ = *µ*_2_*K*_2_ = 100, *K*_1_ = *K*_2_ = 200000.

The impact of wildtype decline rate on the probability of post-change coexistence differs for rescue from SGV and DNM. When rescue occurs from SGV, faster decline of either wildtype promotes the probability of coexistence (Figures 4c) by increasing the rescue probability of each species (Figure 4a,b) despite strengthening survival interference (Figure 4d). The increased interference occurs because faster wildtype decline allows for faster increases in the number of mutants and hence a faster increase in their competitive effect.

When rescue occurs from DNM, faster decline of either wildtype tends to reduce the ultimate probability of coexistence (Figures 4c,e). The effect of wildtype decline on survival interference depends on the symmetry of competition. When competition is symmetric, the strongest interference occurs when wildtype decline rates are highly asymmetric (upper-left and bottom-right corners of Figure 5d). Given it is rescued, the faster declining species will tend to sweep earlier, more strongly inhibiting mutant establishment in the other species.

When competition is highly asymmetric, the strongest interference occurs when the relatively stronger competitor declines quickly and the weaker competitor declines slowly (bottom-right corner of Figure 5f). In this case, rescue of the stronger competitor requires mutants to sweep early, which strongly inhibits the establishment of mutants in the other species, which tend to arrive later. In contrast, when the stronger competitor declines slowly and the weaker competitor declines quickly (upper-left region in Figure 5f), survival interference is weak because mutants of the strong competitor tend to sweep later than mutants of the weaker competitor, at which point the weaker mutants are likely to already be established.

### 3.3 Community rescue

Given the complexity of the analysis with only two species, how are we to scale up to predict the response of realistically large communities? To take a step forward, we start by ignoring the trickiest aspect of the two-species analysis, namely the competition exerted by mutants (and thus survival interference). Then, if we can calculate a probability of rescue for each species, 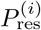, the probability that *k* out of *N* competing species are rescued follows a Poisson binomial distribution, with probability generating function 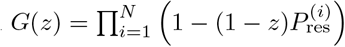. This approach may give a reasonable approximation when competition between mutants in different species is weak and, when compared to simulations, illustrates the strength of survival interference.

In Section S3, we describe how to calculate 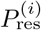 based on our one-species approach. To illustrate the results, we chose parameter values following Van Eldijk et al., 2020, which were inspired by the neutral theory of biodiversity (Hubbell, 1997). In particular, we assume that all *N* species have identical intrinsic growth rates for the wildtypes and rescue mutants, denoted by *r*^wt^ and *r*^mut^, respectively, and share the same wiltype-to-mutant mutation rate, *µ*. At the onset of environmental change, the total community density is *n*_0_, corresponding to an absolute population size of *n*_0_*K*, and the density of the *N* species follows an parameterized distribution function, with the initial fraction of species *i* being *ϕ*_*i*_.

As expected, the approximations closely match simulations (in which mutants do exert competition) when interspecific competition is not too strong (*β* = 0 and 0.5 in Figure 6), as survival interference is then weak, and overestimate simulated rescue probabilities when interspecific is strong (*β* = 1 in Figure 6). In general, initially rarer species have a lower rescue probability (Figure 6a) due to a limited supply of beneficial mutations, even though in the absence of interspecific competition (*β* = 0) their smaller population size means rescue mutants enjoy a higher establishment probability due to weaker intraspecific competition. Given the same initial density of each species, stronger interspecific competition reduces both the expectation and the variation in the number of rescued species (Figures 6b,c) by both reducing the rescue probability of each species (Figure 6a) and strengthening survival interference (Figure 6d).

**Figure 6:**
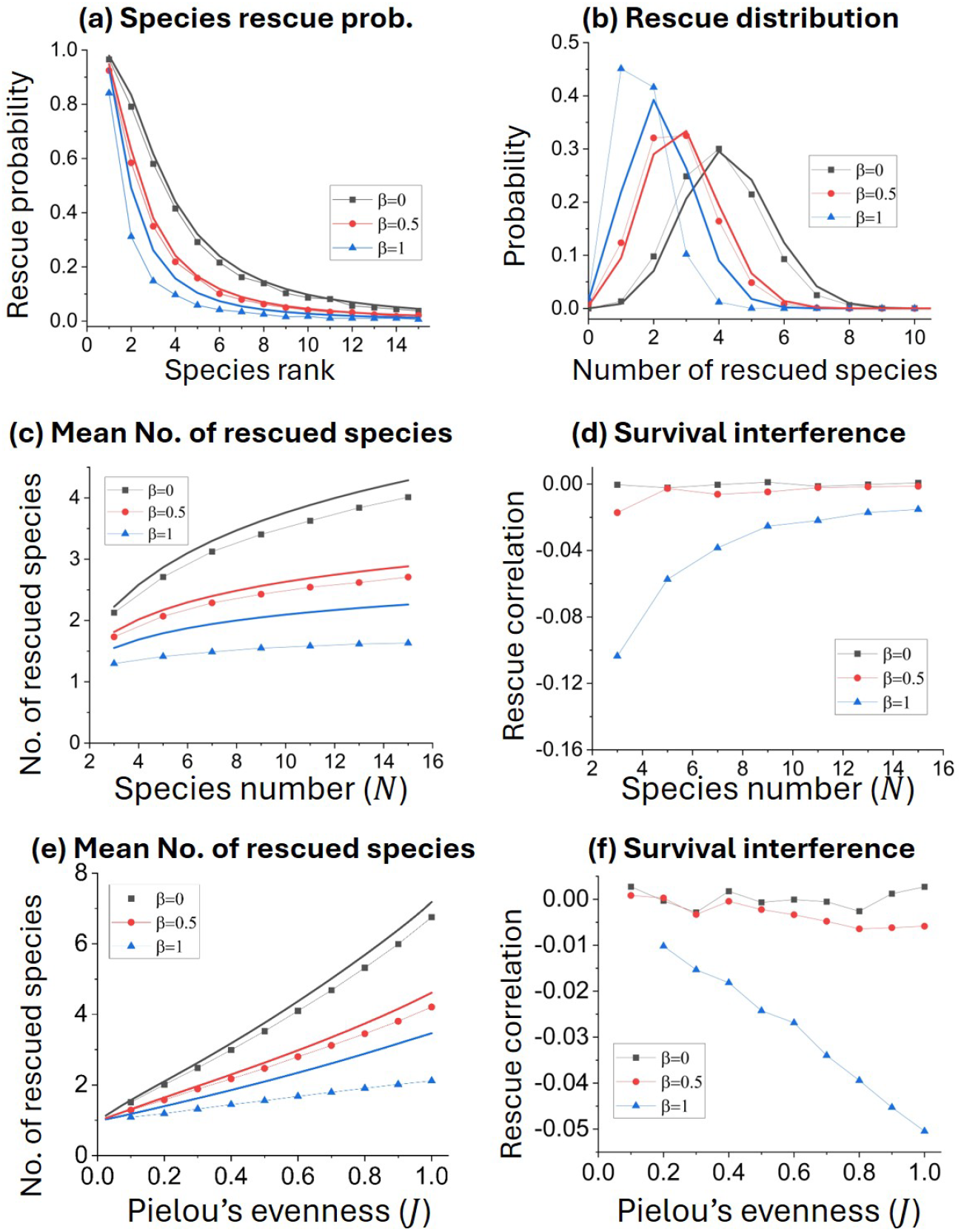
Impacts of interspecific competition strength, species number, and species evenness on community rescue. The figure shows the results for rescue from DNM when when all interspecific competition coefficients, *β*_*ij*_ and 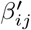, are equal to *β* (results are similar for rescue from SGV; Figure S7). Connected dots are from simulations with 10^5^ replicates and thin lines are analytical approximations (detailed in Section S3), which neglect survival interference. Initial densities follow a power-law rank abundance curve, 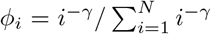. **(a) & (b)** The rescue probability of each species and the probability distribution of the number of rescued species in a community initially with 15 species. **(c) & (d)** Impacts of species number *N*, where we adjust the value of *γ* across *n* such that Pielou’s evenness 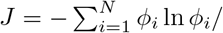 ln *N* (Smith and Wilson, 1996) remains constant. The overall correlation between the rescue outcomes of species is measured by averaging correlation coefficients in rescue events across all pairs of species. **(e) & (f)** Impacts of species evenness (*J*) in a community initially with 10 species. Parameters: *r*^wt^ = − 0.05, *r*^mut^ = 0.1, *n*_0_ = 0.2, *µK* = 20, *K* = 100000, *J* = 0.5.

Given the same initial total density, increasing the number of species leads to a higher expected number of rescued species (but a lower fraction; Figure 6c). This is attributable to both a lower disparity in rescue probabilities among species (resulting in a larger product of rescue probabilities across the same number of species; Figure S8) and, under strong inter-specific competition, reduced survival interference among species (Figure 6d). Intuitively, survival interference can be partially offset by indirect interactions: inhibition from species 1 to species 2 can weaken inhibition exerted by species 2 on species 3. Although indirect interactions are weaker than direct interactions, their number increases more rapidly with species number *N* as each species is involved in only *N* − 1 direct interactions but (*N* − 1)(*N* − 2) second-order indirect interactions. The rescue outcome also depends on the distribution of species densities, *ϕ*_*i*_. Greater evenness in species densities increases the expected number of rescued species, despite causing stronger interference (Figures 6e,f). When we vary both the species number and density evenness, the expected number of species increases nearly linearly with Shannon’s entropy (which incorporates both species number and evenness), 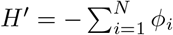 ln *ϕ*_*i*_ (Figure S9). Moreover, the expected number of rescued species increases faster with the number of species when evenness is higher (Figure S9).

## Discussion

The persistence of biodiversity through environmental change will sometimes require evolutionary rescue in one of more species. Whether rescue occurs will depend on interactions between individuals both within and between species. Here we have built, analyzed, and simulated relatively simple stochastic models of evolutionary rescue with competition to improve our understanding of how ecological communities will respond to sudden environmental change.

For within-species competition, we have developed new closed-form approximations to aid intuition. Czuppon et al., 2023 recently found that competition’s impact on the establishment of mutants from the standing genetic variation was determined by the initial density of the wildtype divided by the selective advantage of the mutant, *n*_*a*_(0)*/*(*r*_*A*_ − *r*_*a*_) in our notation. We show that this remains so under non-symmetric competition (*α*_*Aa*_≠ 1, Equation 3) and, naturally, the same term determines the effect of competition on rescue from standing genetic variation (Equation 5). The impact of competition is more complicated for rescue by de novo mutation (Equation 6), but a key factor reducing the probability of rescue is (*n*_*a*_(0) − *r*_*a*_)*/r*_*A*_. Overall then, intraspecific competition will hinder rescue by either SGV or DNM more when the mutant has a smaller intrinsic advantage, *r*_*A*_, and this is therefore when accelerating the wildtype’s decline will lead to the most competitive release. Uecker et al., 2014 numerically found that the probability of rescue by either SGV or DNM was minimized at an intermediate wildtype decline rate when the mutant was not too deleterious prior to change (in our notation, 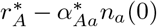 not too negative), so that there was plenty of standing genetic variation, and when competition on the mutant was strong post-change (*α*_*Aa*_ large), so that faster decline of the wildtype greatly increased mutant establishment. Our approximation recapitulates this minimum (Figure 2) and our model formulation, with competition modeled similarly before and after change, emphasizes that this minimum is most likely to occur when the environmental change increases niche overlap between the mutant and wildtype, 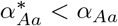, e.g., when there are fewer types of resources to utilize in the new environment.

Between-species competition has three main effects. Similar to within-species competition, competition between wildtypes accelerates wildtype decline and competition from wildtypes reduces mutant establishment. More uniquely, competition also occurs between mutants in different species. In particular, mutant establishment and sequential sweeping in one species reduces the chances a mutant establishes in the other, as previously suggested (Van Eldijk et al., 2020). We call this survival interference in analogy with selective interference (Otto, 2021), in particular competition between two unique mutant lineages that compete for fixation, i.e., the Fisher-Muller effect (as modeled by Gerrish and Lenski, 1998). In both survival and selective interference, beneficial mutations compete with one another and hinder each other’s success. In survival interference, these mutations are in different species and compete for establishment rather than fixation. Both mutations may succeed in establishing, while two mutations cannot fix in one population without recombining onto the same background. For this reason, selective interference is thought to be the key reason sex and recombination have evolved (Otto, 2021). Investigating the analogous effects of introgression and horizontal gene transfer in our multi-species setting would be particularly interesting, as a sweep in a competitor then hinders mutant production and establishment in a focal species but also increases the chances of acquiring a mutant by horizontal transfer (Hildegard Uecker, personal communication).

The persistence of coexistence through environmental change requires mutual invasibility in the new environment. In our model, we have mutual invasibility when *λ < k*_*B*_*/k*_*A*_ *<* 1*/λ*, where 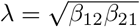 is niche overlap and 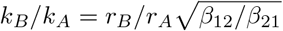 is the mutant competitive ability ratio (Letten et al., 2021). This condition guarantees that a late-arriving mutant has a positive establishment probability. However, it does not guarantee that a mutant establishes, or even ever arises. The persistence of coexistence thus relies not only on mutual invasibility, but on the stochastic arrival and establishment of mutants in both species. Arrival is most likely early, when there are plenty of wildtypes to mutate from, but this is also when the wildtypes suppress establishment most. On the other hand, late-arriving mutants are at risk of being suppressed by competition from mutants in the other species. Once a mutant has established in one species, establishment of the other peaks for a short duration where they have some competitive release from the wildtypes and not yet too much interference from the established mutant (Figure 3). This process is quite similar to community assembly in a sink habitat, where each species may arrive at the habitat at different times. Species that arrive and establish mutants earlier hinder adaptive colonization by later-arriving species, potentially creating strong priority effects through survival interference (Fukami, 2015).

Our model of rescue in a community with an arbitrary number of competing species allows for arbitrary strengths of interspecific competition between wildtypes and wildtypes, *β*_*ij*_, and wildtypes on mutants, 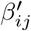. However, when illustrating our approach we only explored the case where all species compete equally (all 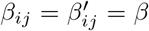). We hope that our approach facilitates future studies that explore the consequences of mutants experiencing different competition than wildtypes (Letten et al., 2021) and investigate how community rescue outcomes depend on characteristics of species interaction networks (e.g., connectivity and modularity; Barabás et al., 2016; Landi et al., 2018; Strydom et al., 2021), potentially with mixtures of competition, predation, and mutualism.

## Acknowledgements

Thanks to Peter Czuppon, Puneeth Deraje, Rebekah Hall, Shaopeng Wang, and Mete Yuksel for feedback. Thanks to the Natural Sciences and Engineering Research Council of Canada (RGPIN–2021-03207 to MMO) and the Department of Ecology and Evolutionary Biology at the University of Toronto (postdoctoral fellowship to KX) for funding.

## Supplementary materials

### S1 One species

#### S1.1 Mutant establishment probability

When a mutant with time-varying birth *b* and death *d* rate arises at time *t*, the probability it establishes is (Eqn 16a in Uecker and Hermisson, 2011)

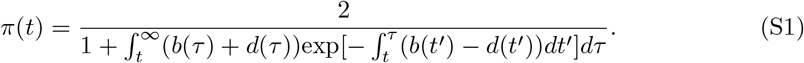

Our model describes individuals in terms of their growth rate, *r*. To use this establishment probability we need to divide net growth into birth and death, *r* = *b* − *d*. We take the discrete time simulations as the truth we wish to approximate. There an individual with growth rate *r* has a Poisson number of offspring with mean *W* = exp(*r*). Following Appendix A in Uecker et al., 2014, when *r* is small the variance in offspring number is nearly 1, requiring *b* + *d* to have the correct rate of drift in the continuous-time approximation. This means the establishment probability is approximately (Eqn A4 in Uecker et al., 2014)

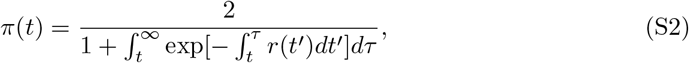

which depends only on the growth rate, *r*.

To obtain the establishment probability of a rare mutant in the one species case, we treat the dynamics of the wildtype deterministically (solving Equation 1 for *n*_*a*_(*t*) with *µ* = 0 and *n*_*A*_ = 0) and substitute the realized growth rate, *r*(*t*) = *r*_*A*_ − *α*_*Aa*_*n*_*a*_(*t*), into Equation S2, giving

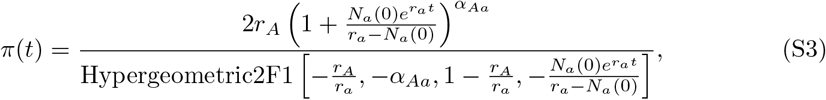

where Hypergeometric2F1[] is the Gaussian hypergeometric function (Abramowitz and Stegun, 1965).

Without competition from the wildtype (*α*_*Aa*_ = 0), Equation S3 reduces to 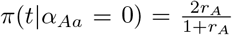 (Uecker et al., 2014). When a wildtype competes with a mutant as strongly as another mutant does, *α*_*Aa*_ = 1, Equation S3 simplifies to

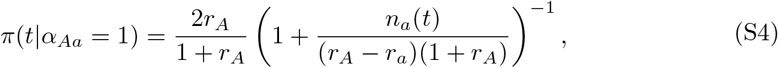

where *n*_*a*_(*t*) is number of wildtypes at time *t* (see Equation 4). Setting *t* = 0, Equation (S4) recovers Equation 3 of Czuppon et al., 2023 for establishment from the standing genetic variation (precise equivalence of their continuous-time analysis and our approximation of our discrete-time model requires their birth and death rates to be 1/2). As time goes to infinity the wildtype goes extinct and *π*(*t*|*α*_*Aa*_ = 1) approaches *π*(*t*|*α*_*Aa*_ = 0).

More importantly, the probability of mutant establishment (Equation S3) declines roughly exponentially with competition strength *α*_*Aa*_, so a useful approximation is a linear interpolation on the log scale between *α*_*Aa*_ = 0 and *α*_*Aa*_ = 1,

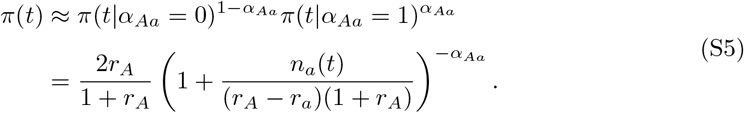

#### S1.2 Rescue from SGV

We assume the population is at equilibrium at the time of environmental change and the wildtype density is constant at *n*_*a*_(0). Mutant density is then determined by a balance between mutational input, *n*_*a*_(0)*µ*^∗^, a negative growth rate when rare, 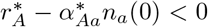, and drift. The probability there are initially *x* mutants is (Orr and Unckless, 2008)

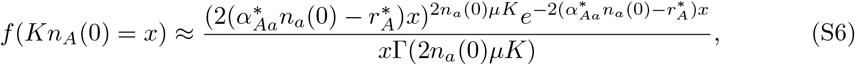

which gives expected initial density 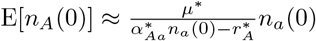 when *µ*^∗^ ≪ 1.

Integrating over the distribution of mutant densities (Equation S6), the probability of rescue from SGV is

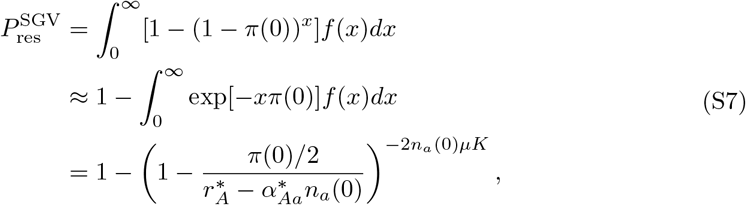

where the approximation assumes *π*(0) ≪ 1.

#### S1.3 Rescue from DNM

The probability of rescue from DNM by time *t* is (Orr and Unckless, 2008)

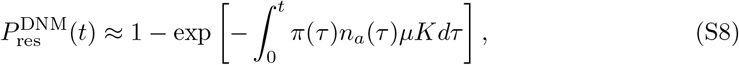

where *n*_*a*_(*τ*)*µK* is the expected number of mutants that occur at time *τ* and the approximation assumes the establishment probability is small (*π*(*τ*) ≪ 1).

Evaluating Equation S8 with Equations 4 and S3 does not result in an analytical solution. Motivated by our approximation of mutant establishment probability (Equation 3), we linearly interpolate 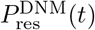 on the log scale between *α*_*Aa*_ = 0 and *α*_*Aa*_ = 1 to get

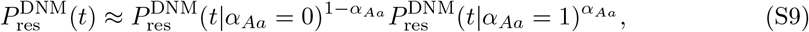

where

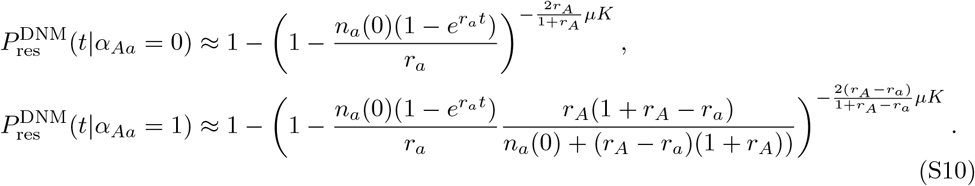

Equation S9 provides a closed-form approximation that applies to any time *t >* 0 and competition strength 0 ≤ *α*_*Aa*_ ≤ 1. The ultimate rescue probabilities are

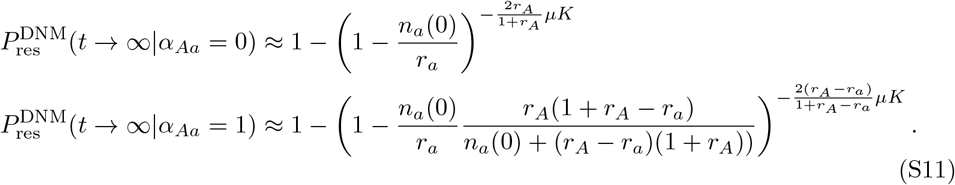

under the weak-growth approximation (*n*_*a*_(0), *r*_*A*_, *r*_*a*_ ≪ 1), these are approximately

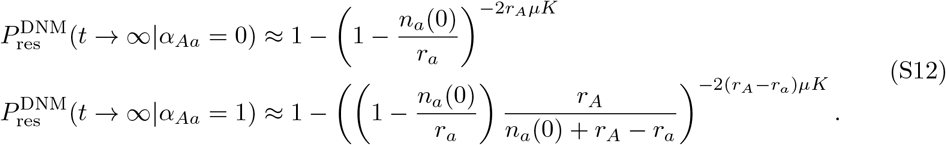

### S2 Two species

#### S2.1 Analytical expressions

In the two species system, we first derive the survival probabilities when stable coexistence between the mutants in the post-change environment is possible, *λ < k*_*B*_*/k*_*A*_ *<* 1*/λ* (Letten et al., 2021). We then discuss how the derivation differs when coexistence is not possible.

Establishment of the rescue allele in a focal species will depend on whether the rescue allele in the other species has established or not. Let 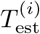 be the time when a mutant lineage that will establish arises in species *i*. Without loss of generality, let the first establishing mutant lineage arise in species 1. The number of pre-existing copies of the mutant allele, *A*, is 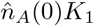 and the number of *de novo* mutants arising at time *t* is *µ*_1_*n*_*a*_(*t*)*K*_1_. The probability that a mutant lineage that will establish arises by time *t* is 1 minus the probability that all pre-existing and new mutations by time *t* do not establish,

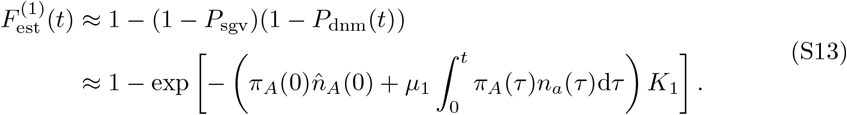

The establishment probability of each mutant allele that exists at time *t, π*_*A*_(*t*), is given by Equation S1 with growth rate 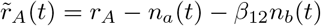 and deterministic wildtype dynamics.

Assuming instead the first establishing mutant arises in species 2, a change of subscripts gives 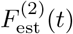.

Both species will go extinct if rescue alleles fail to establish in both species, which occurs with probability

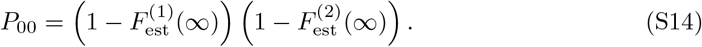

Deriving the probability that one species survives but the other goes extinct, *P*_01_ and *P*_10_, is more complicated. The probability that a rescue allele establishes in species 1 first, between time *t* and *t* + d*t*, is

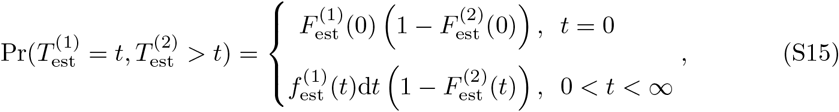

where 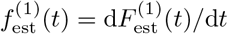. Conditioned on this, the probability that a mutant lineage will never establish in species 2, and thus species 2 goes extinct, is

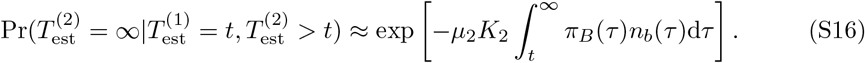

The establishment probability of a mutant lineage in species 2, *π*_*B*_(*t*), can be obtained based on Equation S1 with growth rate 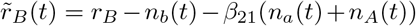, which accounts for interference from mutants in species 1, *A*, and we treat the dynamics of the three strains, *n*_*a*_(*t*), *n*_*b*_(*t*) and *n*_*A*_(*t*) deterministically. The probability that species 1 survives but species 2 goes extinct is thus

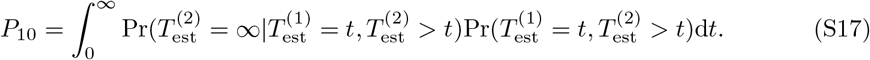

The probability that species 2 survives but species 1 goes extinct, *P*_01_, can be obtained in a similar way. The probability that both species will survive and coexist is *P*_11_ = 1 − *P*_10_ − *P*_01_ − *P*_00_.

When coexistence is not possible in the new environment we need to slightly modify our derivations. Without loss of generality, consider the case where species 2 is competitively excluded given species 1 establishes, but not vice versa. The calculation of *P*_00_ and *P*_01_ are unchanged because species 1 goes extinct and thus cannot exclude species 2. However, now *P*_11_ = 0 and the only way species 2 can survive is for species 1 to go extinct, *P*_10_ = 1 − *P*_00_ − *P*_01_. When species 2 competitively excludes species 1 but not vice-versa we similarly recalculate *P*_01_ as 1 − *P*_00_ − *P*_10_. When the two species exclude each other in the new environment (i.e., *r*_*A*_ − *β*_12_*r*_*B*_ *<* 0, *r*_*B*_ − *β*_21_*r*_*A*_ *<* 0), the probability that a species survives is the probability it has a mutant that sweeps first and the calculation of *P*_10_ (similarly for *P*_01_) can be simplified to 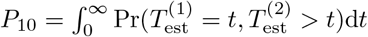.

#### S2.2 Numerical solutions

Due to the complicated dynamics of a two-species system, closed-form solutions for the probability of the four possible outcomes are not available. Even exact numerical solutions are difficult because of several multi-dimensional numeric integrations. We therefore use multiple approximations to get analytical expressions for intermediate steps to speed up numeric calculations.

Before either rescue mutant establishes, the dynamics of the two wild types can be approximated using a recursive solution method. Specifically, we first solve for the density dynamics under intraspecific competition alone (ignore interspecific competition). We then substitute these solutions into the right-hand sides of the differential equations that include interspecific competition (Equation 2) and solve again for the density dynamics (see Mathematica notebook), giving

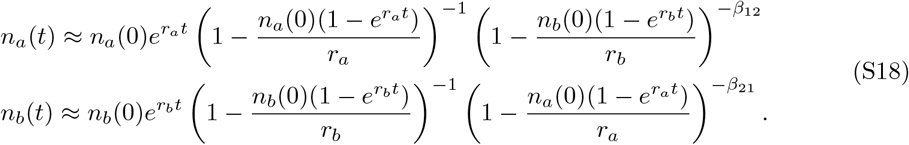

These approximations are used to calculate the establishment probability of the mutants before any start to sweep (Equation S1) and the time until the first mutant starts sweeping (Equation S13).

We then obtain an approximate analytical solution for the establishment probability of a rescue mutant in one species, conditioned on the rescue mutant in the other species having started to sweep first. Without loss of generality, we focus on the establishment probability of allele *B*, conditioned on allele *A* sweeping first at time *t*_1_, denoted by 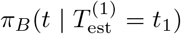. For analytical tractability, after allele *A* begins to sweep, we approximate the dynamics of the three alleles (*a, b*, and *A*) by completely neglecting competition between alleles, which will overestimate their densities and thus underestimate the establishment probability of allele *B*. Under this assumption, the three strains grow logistically as

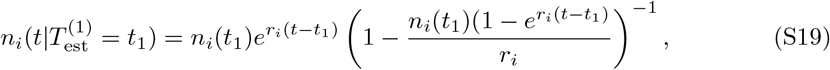

where *n*_*i*_(*t*_1_) is the density of allele *i* at the time when allele *A* starts to sweep. The density of wildtypes at time *t*_1_ are obtained from Equation S18 and the initial density of the mutant lineage that establishes at time *t*_1_ is effectively *n*_*A*_(*t*_1_) = 1*/*(*K*_1_*π*_*A*_(*t*_1_)) (Orr and Unckless, 2008), where *π*_*A*_(*t*_1_) is the establishment probability of the mutant before any other mutants have established. The establishment probability of allele *B*, 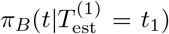, can then be calculated using Equation S1. However, a closed-form solution is only available when *β*_21_ = 0 and *β*_21_ = 1. We therefore made additional approximations to obtain an analytical solution for general *β*_21_. Note that the establishment probability is a function of the realized growth rate, 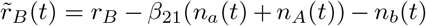. Since an analytical solution is available when *β*_21_ = 1, we consider a hypothetical system with 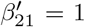 and densities 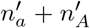 that give approximately the same amount of competition, 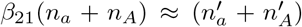 (see Figure S10 for illustration). To do this we give allele *a* initial density 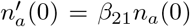, so that 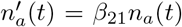 for all *t*. For allele *A*, we give it growth rate 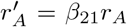 to ensure that the realized growth rate of allele *B* is the same when allele *A* reaches carrying capacity, 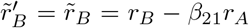. The initial density of allele *A* is set to 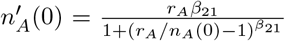, such that the inflection point of the logistic growth (i.e., the time at which 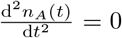) is the same for both populations (intersection of the two lines in Figure S10a).

### S3 Community rescue

To extend to *N* species we ignore competition exerted by mutants so that the rescue of each species is independent. The probability that *k* out *N* species are rescued then follows a Poisson binomial distribution, with probability generating function 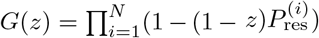 and moment generating function 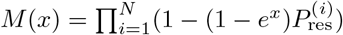, where 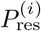 is the probability that species *i* is rescued. The probability that *k* out *N* species are rescued is 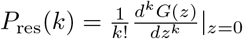 and the expected number of rescued species is 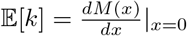.

As in the two-species model, we scale within-species competition to 1. We denote competition from the wildtype in species *j* to the mutant in species *i* as *β*_*ij*_ and competition from the wildtype in species *j* to the wildtype in species *i* as 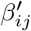. The intrinsic growth rates of the wildtype and mutant in species *i* are 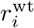 and 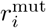, respectively, and the mutation rate from the wildtype to the mutant allele is *µ*_*i*_. The density of wildtypes in species *i* at time *t* is 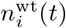.

Rescue probabilities, 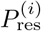, are calculated as follows. We first solve for the wildtype dynamics, 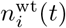, of all *N* species numerically. Then we obtain the establishment probability of rescue mutants in each species based on Equation S2 with the realized growth rate 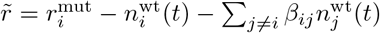. The probability that species *i* is rescued from DNM is then given by Equation S8 with wildtype density 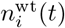 and mutation rate *µ*_*i*_. For rescue from SGV, instead of dynamically modeling densities prior to environmental change, we assume the initial wildtype density of species *i* is fixed at 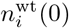. The probability distribution of the initial number of mutants and the probability of rescue from SGV are then given by Equations S6 and S7, respectively, replacing the realized growth rate, 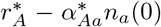, by the pre-change selection coefficient, 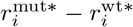.

We can also derive closed-form expressions for the probability of rescue in special cases. Without interspecific competition (all 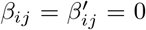), the probability of rescue from DNM is given by the first row of Equation S11 with initial wildtype density 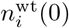. For rescue from SGV, the rescue probability is given by Equation 5 with the realized growth rate, 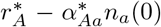, replaced by the pre-change selection coefficient, 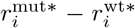, a wildtype density of 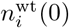, and establishment probability given by Equation S4 with wildtype density 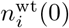.

When all individuals compete equally, 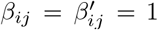, and the wildtype growth rates are identical across all *N* species, 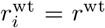, the total wildtype density of the community, 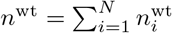, declines logistically and the relative abundance of each species, *ϕ*_*i*_, remains constant. In this case, the probability that species *i* is rescued from DNM is given by the second row of Equation S11, with the initial wildtype density replaced by the total community density, *n*^wt^(0), and the mutation rate replaced by *µ*_*i*_*ϕ*_*i*_. For rescue from SGV, the rescue probability is calculated as described in the preceding paragraph, with establishment probability given by Equation S4 with the wildtype density *n*^wt^(0).

## Supplementary Figures

**Figure S1:**
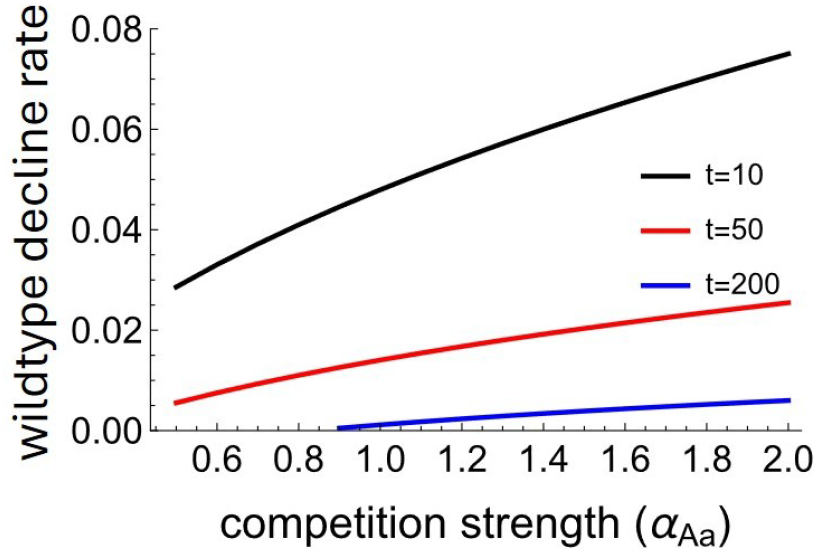
Impact of intraspecific competition strength on the wildtype decline rate that maximizes the probability of rescue from DNM. Results are obtained from S9. Parameters: 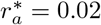, *µK* = 20, *r*_*A*_ = 0.01.

**Figure S2:**
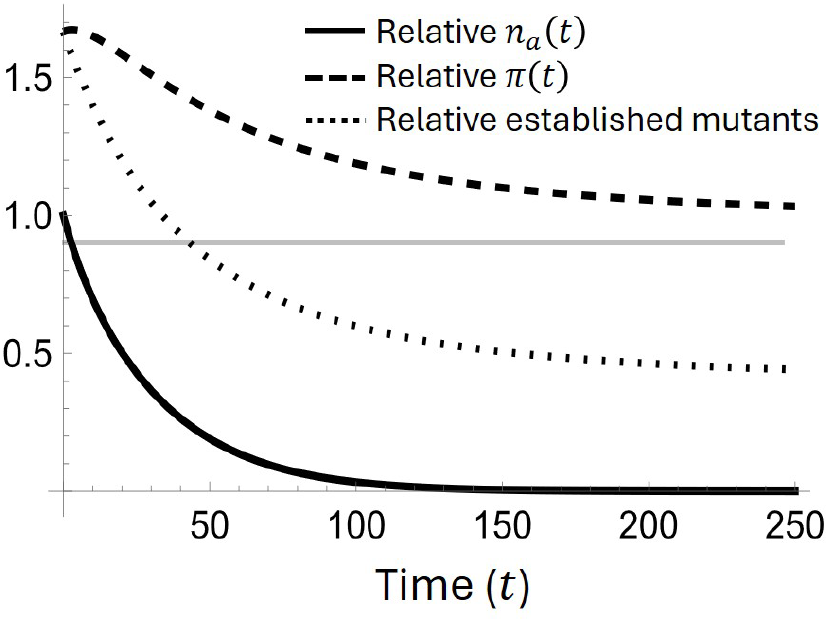
The costs and benefits of wildtype decline. The figure shows the expected number of mutations *µn*_a_ (*t*)*K* (solid), the establishment probability *π*(*t*) (dashed), and the expected cumulative number of established mutants 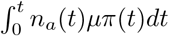 (dotted) under fast wildtype decline (*r*_*a*_ = −0.05) relative to that under slow wildtype decline (*r*_*a*_ = −0.01). Results are calculated based on Equation 4 for *n*_*a*_(*t*) and Equation S3 for *π*(*t*). Parameters are the same as those in Figure 2d.

**Figure S3:**
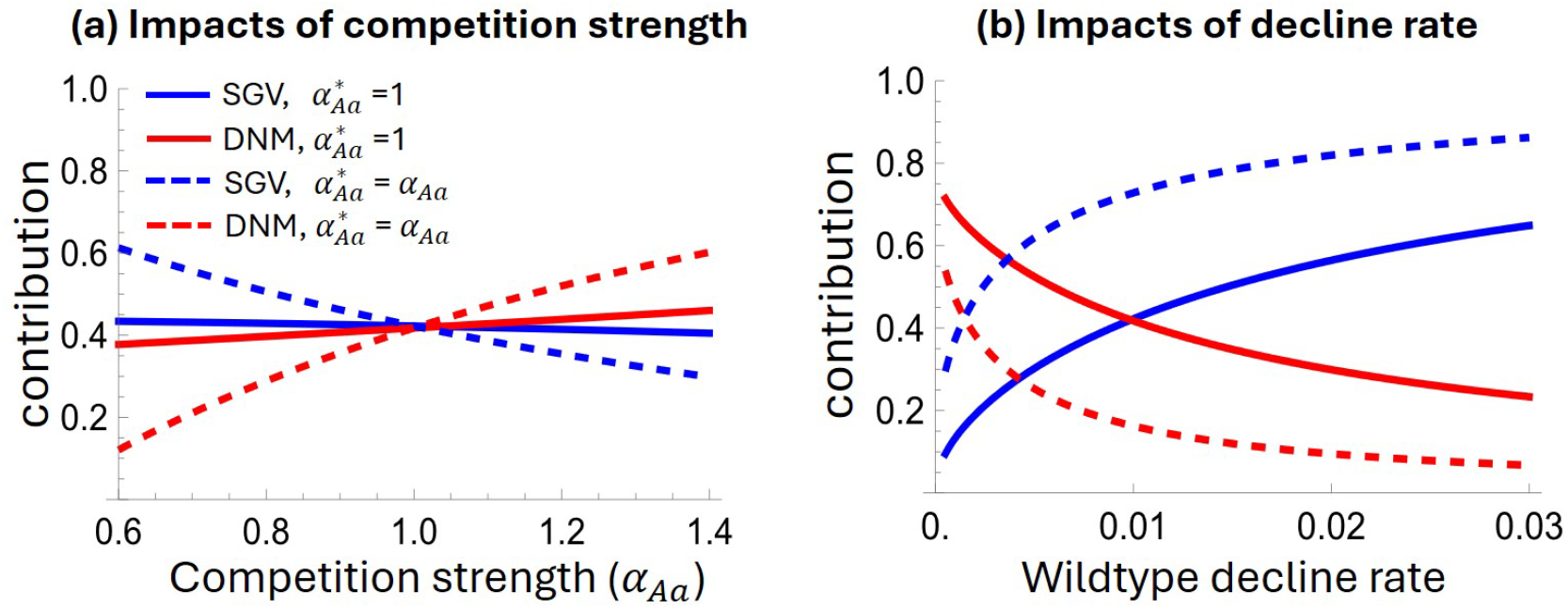
Impacts of intraspecific competition strength and wildtype decline rate on the relative contributions from SGV and DNM to rescue. The relative contributions are calculated as 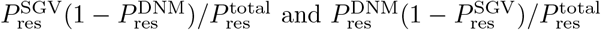 Parameters in panels (a) and (b) are the same as those used in panels (e) and (f) of Figure 2, respectively. Results are calculated based on Equation 5 for 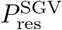 and Equation 6 for 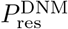.

**Figure S4:**
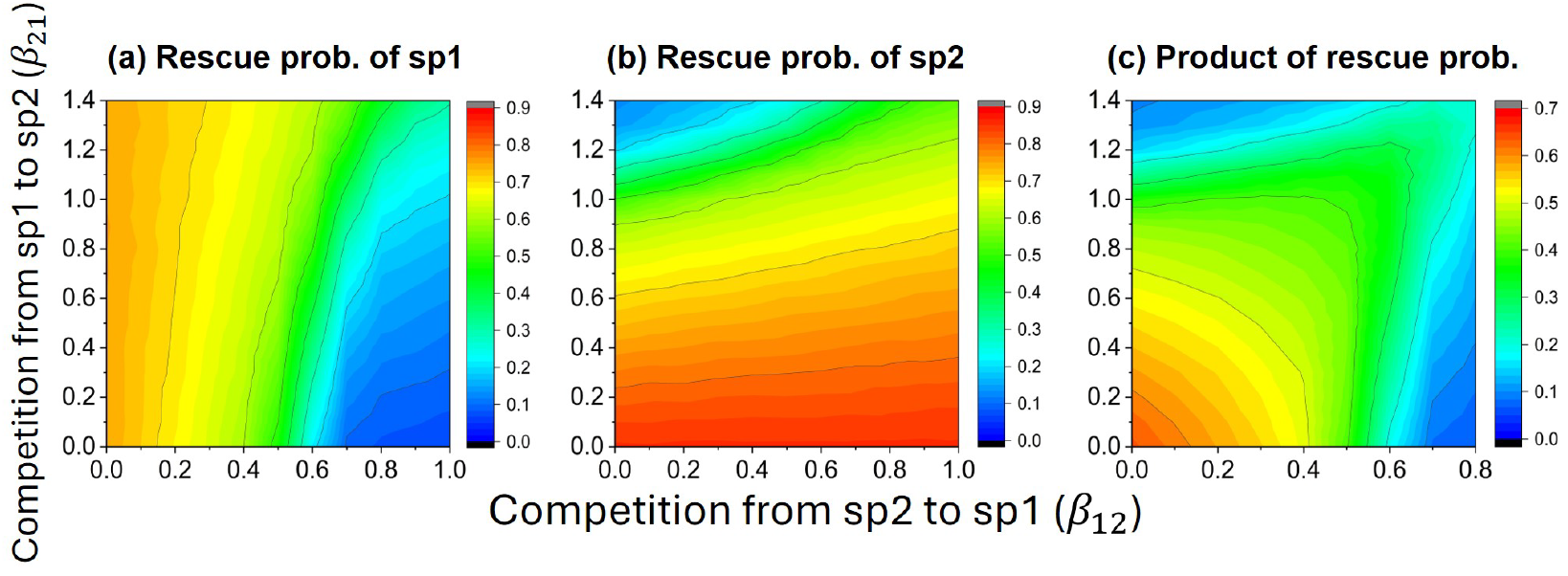
Impact of interspecific competition strength on the rescue probability of each species when rescue occurs from DNM. Panel (c) shows the product of the rescue probabilities of species 1 and 2, demonstrating that the reduced joint rescue probability (Figure 4c) is due to both a decreased rescue probability in one species and stronger competitive interference (Figure 4d). Results are from simulations with 10^5^ replicates. Parameters the same as those in Figure 4.

**Figure S5:**
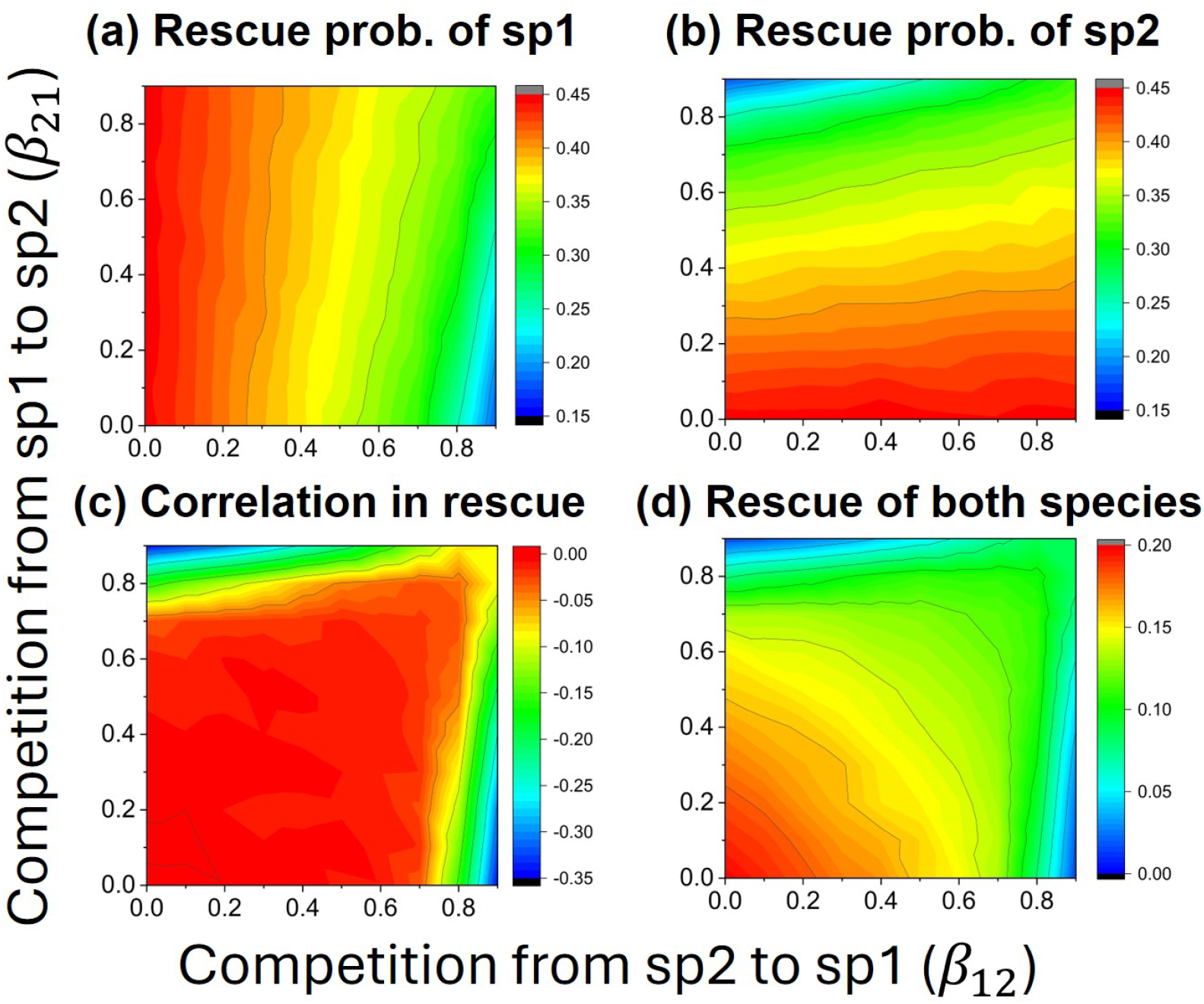
Impacts of interspecific competition strength on rescue from SGV in a two-species system. Results are from simulations with 10^5^ replicates. The initial densities of the two species are fixed to be *n*_1_(0) = *n*_2_(0) = 0.03. Other parameters: 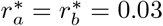, *r*_*a*_ = *r*_*b*_ = −0.01, 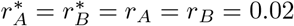, *µ*_1_*K*_1_ = *µ*_2_*K*_2_ = 10, *K*_1_ = *K*_2_ = 200000.

**Figure S6:**
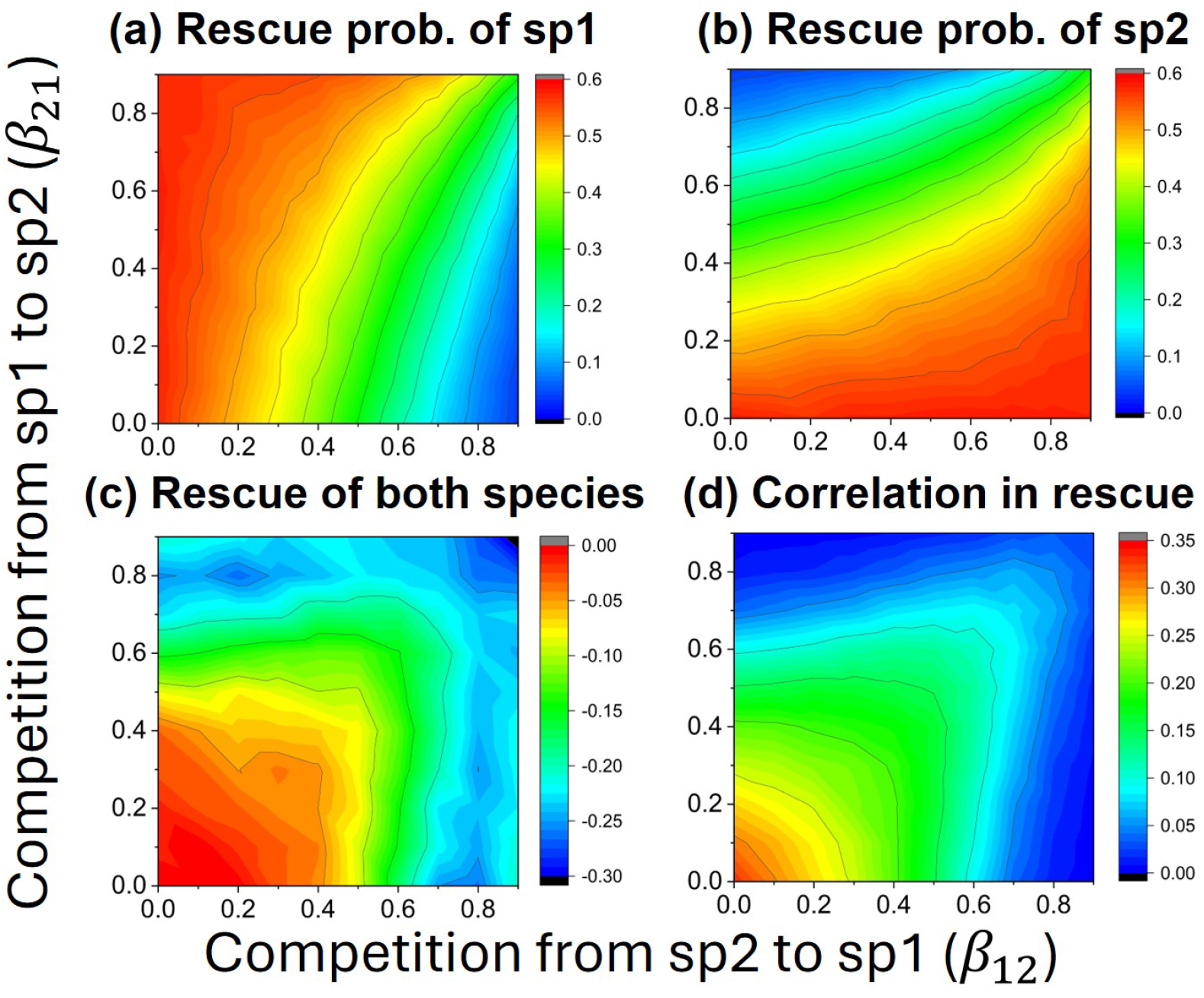
Impacts of interspecific competition strength on rescue from DNM in a two-species system. The results assume competition strengths before and after the environmental shift are identical and the initial densities change with competition strength. Results are from simulations with 10^5^ replicates. Parameters: 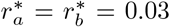, *r*_*a*_ = *r*_*b*_ = −0.01, *r*_*A*_ = *r*_*B*_ = 0.01, *µ*_1_*K*_1_ = *µ*_2_*K*_2_ = 100.

**Figure S7:**
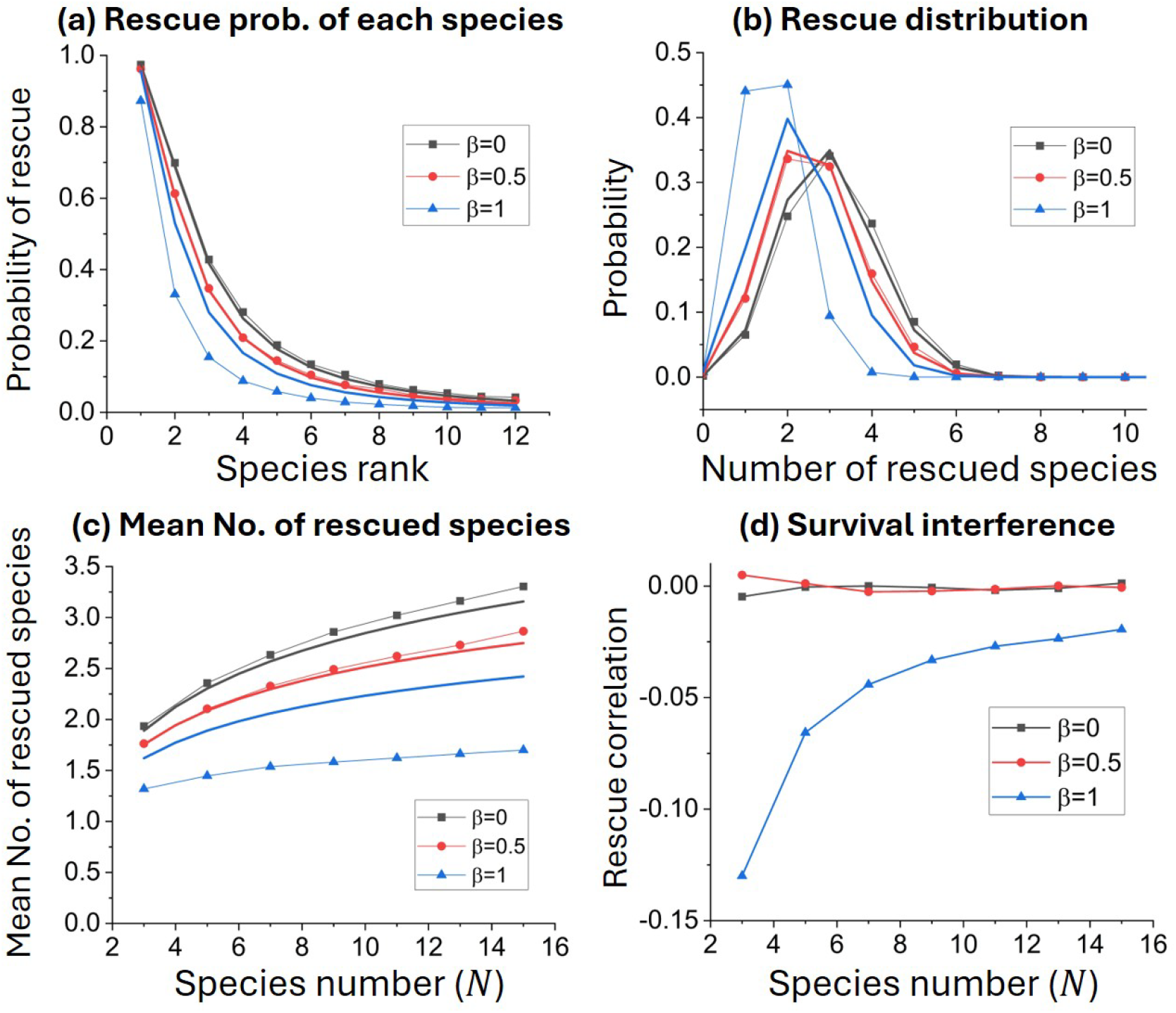
Impacts of interspecific competition strength and species number on community rescue from SGV. The species are equivalent except for their initial densities, which are fixed and follow a power-law rank abundance curve, 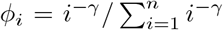. The rescue mutants are assumed to be at selection-mutation-drift balance before the environmental shift. Connected dots are from simulations with 10^5^ replicates and thick lines are analytical approximations (described in Section S3). **(a**,**b)** The rescue probability of each species and the probability distribution of the number of rescued species in a community initially with 12 species. **(c**,**d)** Impact of species number *N* on the mean number of rescued species and survival interference, where the value of *γ* is adjusted with *N* such that Pielou’s evenness 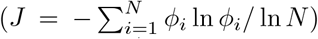 is constant. Other parameters: 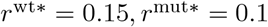, *r*^wt^ = −0.05, *r*^mut^ = 0.1, *n*_0_ = 0.2, *µK* = 20, *K* = 100000, *J* = 0.5.

**Figure S8:**
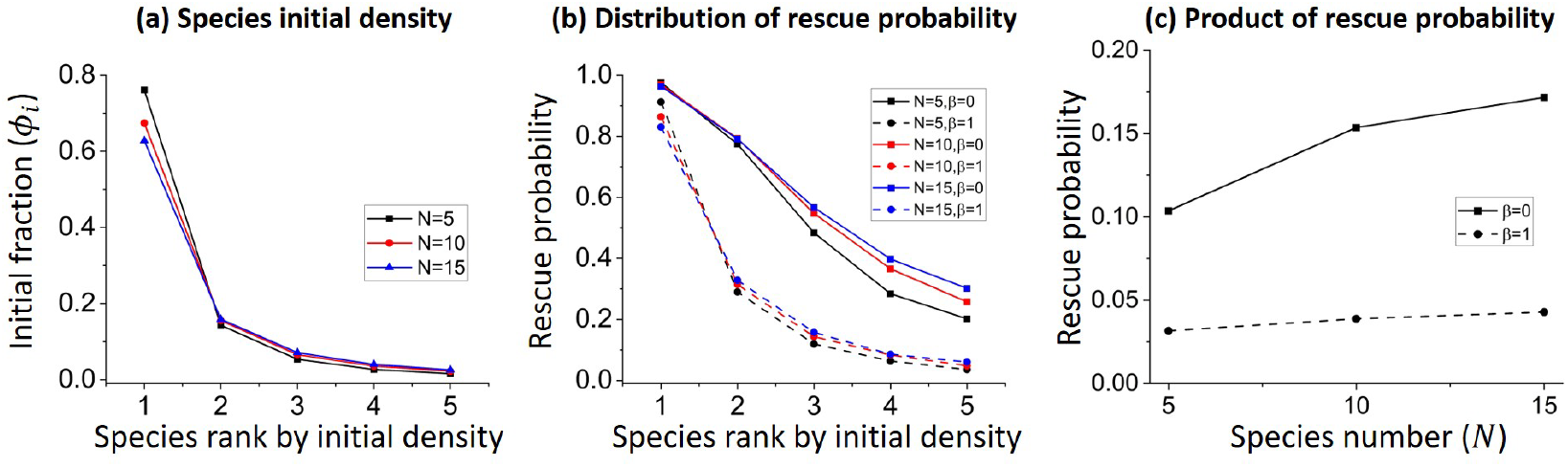
Impacts of species number on community rescue. Results are from 10,000 replicate individual-based simulations of rescue from DNM. **(a)** Effect of species number *N* on the initial densities of the five most abundant species, with the total initial density fixed at *n*_0_ = 0.2 and Pielou’s evenness of the density distribution fixed at *J* = 0.5. **(b)** Effects of species number on the rescue probabilities of the five most abundant species under two interspecific competition strengths. **(c)** Product of the rescue probabilities of species with individual rescue probabilities greater than 0.05 (five species for interspecific competition strength *β* = 0 and four species for *β* = 1). Other parameters are the same as those in Figure 6.

**Figure S9:**
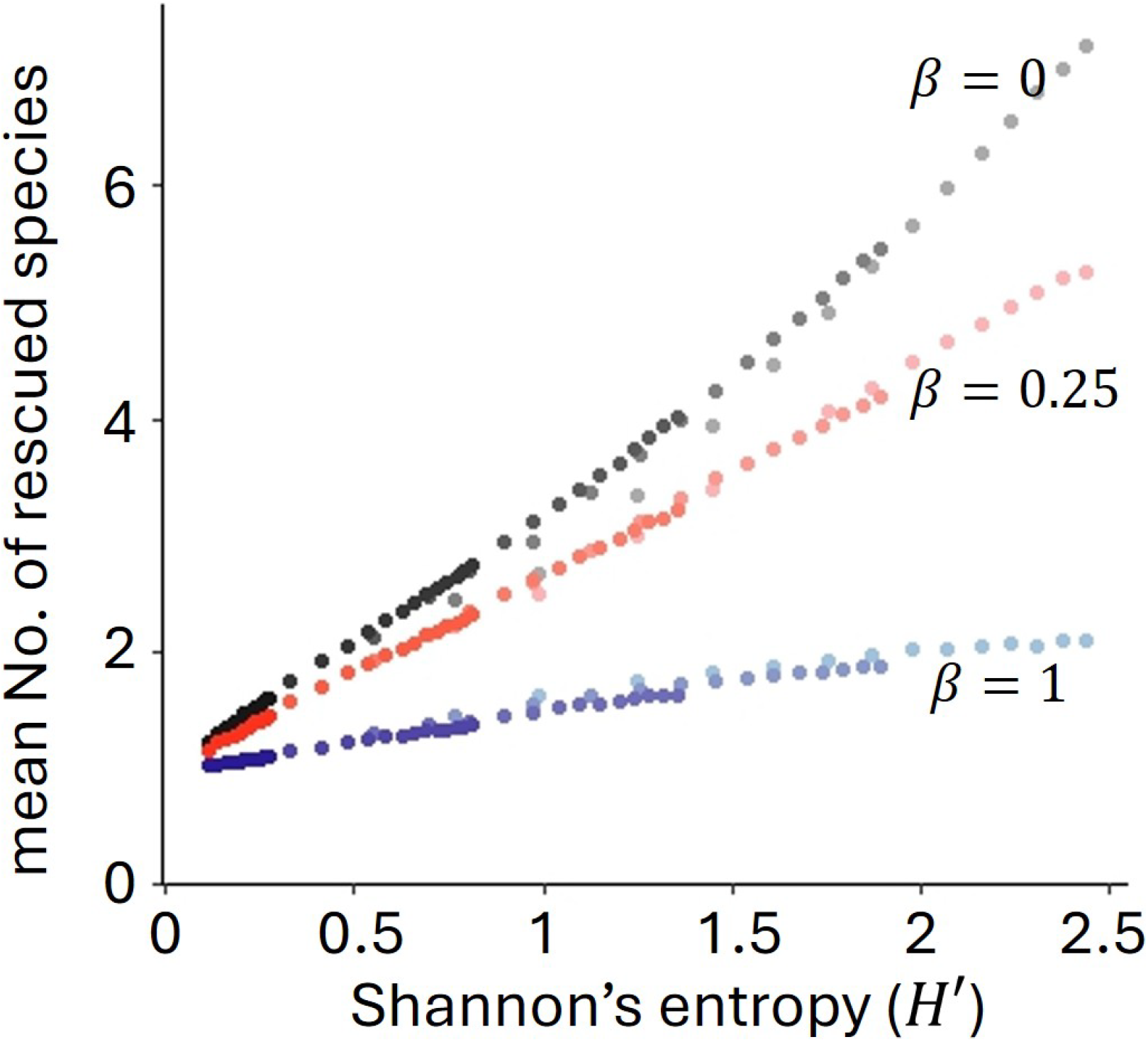
Impacts of Shannon’s entropy on community rescue. Results are from 10,000 replicate simulations of rescue from DNM. The initial species density distribution follows a power-law, *ϕ*_*i*_ ∝ *i*^−*γ*^. Shannon’s entropy is 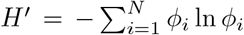. For a given competition strength, *β*, each dot represents one combination of Pielou’s evenness *J* and species number *N*, with all combinations of *J* = 0.1, 0.3, 0.5, 0.7, 0.9 and *N* = 3, 4, …, 15 simulated. The darkness of the dots indicate *J* (becoming progressively lighter as *J* varies from 0.1 to 0.9). Other parameters: *r*^wt^ = − 0.05, *r*^mut^ = 0.1, *n*_0_ = 0.2, *µK* = 20, *K* = 100000.

**Figure S10:**
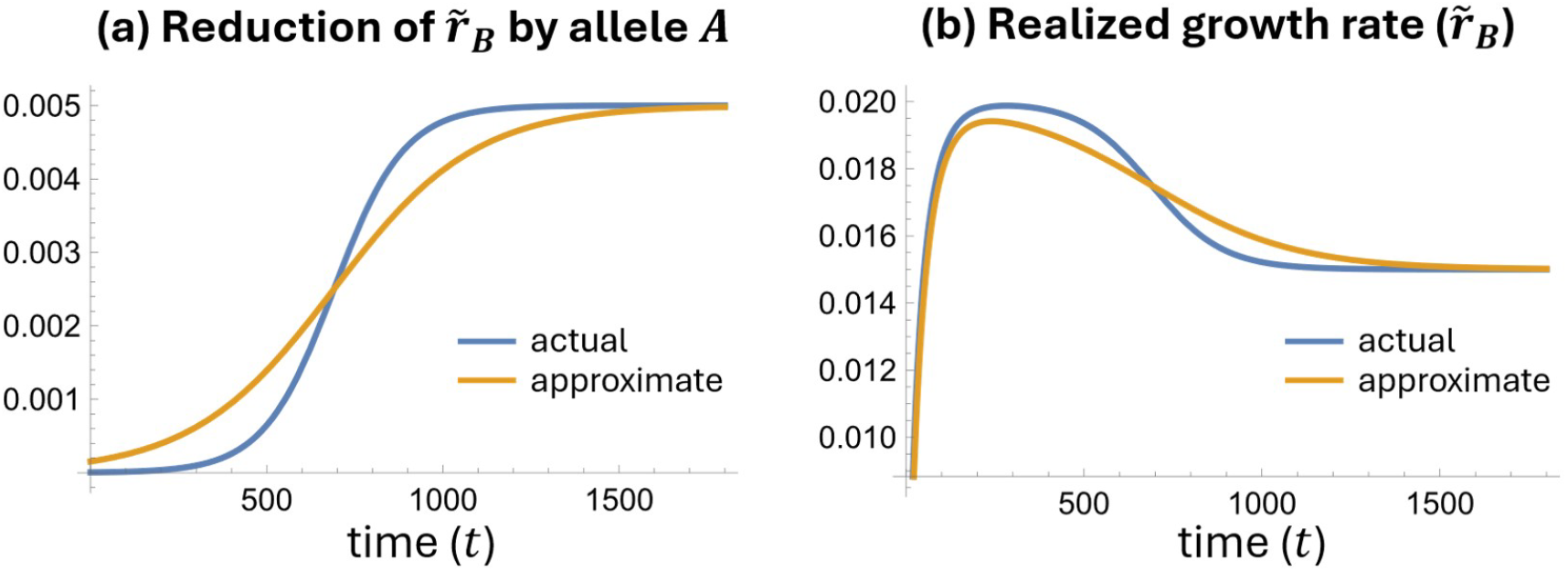
Illustration of the approximation method described in Section S2. **(a)** The reduction of the realized growth rate of allele *B* caused by sweeping of the mutant allele *A, β*_21_*n*_*A*_. **(b)** The realized growth rate of allele *B*, 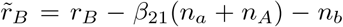. The blue curve gives the actual dynamics (where *β*_21_ = 0.5) and the orange curve shows the dynamics in a hypothetical population with *β*_21_ = 1 and adjusted initial density and growth rates. Parameters: *n*_*a*_(0) = 0.02, *n*_*b*_(0) = 0.01, *n*_*A*_(0) = 0.00001, *r*_*a*_ = *r*_*b*_ = −0.02, *r*_*A*_ = 0.01, *r*_*B*_ = 0.02.

## References

Barabás, G., Michalska-Smith, M. J., & Allesina, S. (2016). The effect of intra-and interspecific competition on coexistence in multispecies communities. The American Naturalist, 188 (1), E1–E12.

Barrett, R. D., & Schluter, D. (2008). Adaptation from standing genetic variation. Trends in Ecology & Evolution, 23 (1), 38–44.

Bell, G. (2017). Evolutionary rescue. Annual Review of Ecology, Evolution, and Systematics, 48, 605–627.

Bomblies, K., & Peichel, C. L. (2022). Genetics of adaptation. Proceedings of the National Academy of Sciences, 119 (30), e2122152119.

Chevin, L., & Lande, R. (2010). When do adaptive plasticity and genetic evolution prevent extinction of a density-regulated population? Evolution, 64 (4), 1143–1150.

Czuppon, P., Day, T., Débarre, F., & Blanquart, F. (2023). A stochastic analysis of the interplay between antibiotic dose, mode of action, and bacterial competition in the evolution of antibiotic resistance. PLoS Computational Biology, 19 (8), e1011364.

Day, T., & Read, A. F. (2016). Does high-dose antimicrobial chemotherapy prevent the evolution of resistance? PLoS computational biology, 12 (1), e1004689.

Délye, C., Jasieniuk, M., & Le Corre, V. (2013). Deciphering the evolution of herbicide resistance in weeds. Trends in Genetics, 29 (11), 649–658.

de Mazancourt, C., Johnson, E., & Barraclough, T. G. (2008). Biodiversity inhibits species’ evolutionary responses to changing environments. Ecology Letters, 11 (4), 380–388.

Fugère, V., Hébert, M., Da Costa, N., Xu, C., Barrett, R., Beisner, B., Bell, G., Fussmann, G., Shapiro, B., Yargeau, V., & Gonzalez, A. (2020). Community rescue in experimental phytoplankton communities facing severe herbicide pollution. Nature Ecology and Evolution, 4 (4), 578–588.

Fukami, T. (2015). Historical contingency in community assembly: Integrating niches, species pools, and priority effects. Annual Review of Ecology, Evolution, and Systematics, 46, 1–23.

Gerrish, P. J., & Lenski, R. E. (1998). The fate of competing beneficial mutations in an asexual population. Genetica, 102 (0), 127–144.

Gomulkiewicz, R., & Holt, R. D. (1995). When does evolution by natural selection prevent extinction? Evolution, 49 (1), 201–207.

Hawkins, N. J., Bass, C., Dixon, A., & Neve, P. (2019). The evolutionary origins of pesticide resistance. Biological Reviews, 94 (1), 135–155.

Hubbell, S. P. (1997). A unified theory of biogeography and relative species abundance and its application to tropical rain forests and coral reefs. Coral reefs, 16 (Suppl 1), S9–S21.

Johansson, J. (2008). Evolutionary responses to environmental changes: How does competition affect adaptation? Evolution, 62 (2), 421–435. 10.1111/j.1558-5646.2007.00301.x

Jones, A. G. (2008). A theoretical quantitative genetic study of negative ecological interactions and extinction times in changing environments. BMC Evolutionary Biology, 8, 119. 10.1186/1471-2148-8-119

Kersten, S., Chang, J., Huber, C. D., Voichek, Y., Lanz, C., Hagmaier, T., Lang, P., Lutz, U., Hirschberg, I., Lerchl, J., & Porri, A. (2023). Standing genetic variation fuels rapid evolution of herbicide resistance in blackgrass. Proceedings of the National Academy of Sciences, 120 (16), e2206808120.

Kivikoski, M., Cairns, J., Hogle, S. L., Pausio, S., Becks, L., Mustonen, V., & Hiltunen, T. (2026). Evolution induced state shifts in a long-term microbial community experiment. Proceedings of the National Academy of Sciences, 123 (22), e2533269123.

Landi, H. O., P. ajd Minoarivelo, Brännström, Å., Hui, C., & Dieckmann, U. (2018). Complexity and stability of ecological networks: A review of the theory. Population Ecology, 60 (4), 319–345.

Letten, A. D., Hall, A. R., & Levine, J. M. (2021). Using ecological coexistence theory to understand antibiotic resistance and microbial competition. Nature Ecology and Evolution, 5 (4), 431–441.

Low-Décarie, E., Kolber, M., Homme, P., Lofano, A., Dumbrell, A., Gonzalez, A., & Bell, G. (2015). Community rescue in experimental metacommunities. Proceedings of the National Academy of Sciences, 112 (46), 14307–14312.

Mallet, J. (2012). The struggle for existence. how the notion of carrying capacity, k, obscures the links between demography, darwinian evolution and speciation. Evolutionary Ecology Research, 14, 627–665.

Nordstrom, S. W., Hufbauer, R. A., Olazcuaga, L., Durkee, L. F., & Melbourne, B. A. (2023). How density dependence, genetic erosion and the extinction vortex impact evolutionary rescue. Proceedings of the Royal Society B: Biological Sciences, 290 (2011).

Orr, H. A., & Unckless, R. L. (2008). Population extinction and the genetics of adaptation. The American Naturalist, 172 (2), 160–169.

Osmond, M. M., & de Mazancourt, C. (2013). How competition affects evolutionary rescue. Philosophical Transactions of the Royal Society B: Biological Sciences, 368 (1610), 20120085. 10.1098/rstb.2012.0085

Otto, S. P. (2021). Selective interference and the evolution of sex. Journal of Heredity, 112 (1), 9–18.

Read, A. F., Day, T., & Huijben, S. (2011). The evolution of drug resistance and the curious orthodoxy of aggressive chemotherapy. Proceedings of the National Academy of Sciences, 108 (supplement 2), 10871–10877.

Smith, B., & Wilson, J. B. (1996). A consumer’s guide to evenness indices. Oikos, 76 (1), 70–82.

Strydom, T., Catchen, M. D., Banville, F., Caron, D., Dansereau, G., Desjardins-Proulx, P. R. F.-M. N., Higino, G., Mercier, B., Gonzalez, A., & Poisot, T. (2021). A roadmap towards predicting species interaction networks (across space and time). Philosophical Transactions of the Royal Society B: Biological Sciences, 367 (1837), 20210063.

Uecker, H., & Hermisson, J. (2011). On the fixation process of a beneficial mutation in a variable environment. Genetics, 188 (4), 915–930.

Uecker, H., Otto, S. P., & Hermisson, J. (2014). Evolutionary rescue in structured populations. The American Naturalist, 183 (1), E17–E35.

Van Eldijk, T. J., Bisschop, K., & Etienne, R. S. (2020). Uniting community ecology and evolutionary rescue theory: Community-wide rescue leads to a rapid loss of rare species. Frontiers in Ecology and Evolution, 8, 552268.

Wilson, B. A., Pennings, P. S., & Petrov, D. A. (2017). Soft selective sweeps in evolutionary rescue. Genetics, 205 (4), 1573–1586.

